# Parallel pathways carrying direction and orientation selective retinal signals to layer 4 of mouse visual cortex

**DOI:** 10.1101/2023.09.18.558281

**Authors:** Helen Wang, Oyshi Dey, Willian N. Lagos, Noor Behnam, Edward M. Callaway, Benjamin K. Stafford

## Abstract

Parallel functional and anatomical visual pathways from the retina to primary visual cortex (V1) via the lateral geniculate nucleus (LGN) are common to many mammalian species, including mice, carnivores and primates. However, the much larger number of retinal ganglion cell (RGC) types that project to the LGN, as well as the more limited lamination of both the LGN and the thalamocortical-recipient layer 4 (L4) in mice, leaves considerable uncertainty about which visual features present in both retina and V1 might be inherited from parallel pathways versus extracted by V1 circuits in the mouse visual system. Here, we explored the relationships between functional properties of L4 V1 neurons and their RGC inputs by taking advantage of two Cre-expressing mouse lines — Nr5a1-Cre and Scnn1a-Tg3-Cre — that each label functionally and anatomically distinct populations of L4 neurons. Visual tuning properties of L4 V1 neurons were evaluated using Cre-dependent expression of GCaMP6s followed by 2-photon calcium imaging. RGCs providing input to these neurons (via LGN) were labeled and characterized using Cre-dependent trans-synaptic retrograde labeling with G-deleted rabies virus. We find significant differences in the tuning of Nr5a1-Cre versus Scnn1a-Tg3-Cre neurons for direction, orientation, spatial frequency, temporal frequency, and speed. Strikingly, a subset of the RGCs had tuning properties that matched the direction and orientation tuning properties of the L4 V1 neurons to which they provided input. Altogether, these results suggest that direction and orientation tuning of V1 neurons may be at least partly inherited from parallel pathways originating in the retina.

## Introduction

The neural circuits that subserve vision originate in the retina where visual information is conveyed by retinal ganglion cells (RGCs) first to the dorsal lateral geniculate nucleus (dLGN) of the thalamus, then to neurons in specific layers of the primary visual cortex (V1). Information is integrated at various stages of this pathway to generate complex response properties in neurons throughout the visual pathway that help give rise to visual perception and behavior (Douglas and Martin, 2004). In the mouse visual system, there are numerous types of cells that encode information in specific ways at each step of the visual pathway. For instance, in the retina there are 40-45 different types of RGCs, each of which are ‘tuned’ to respond preferentially to specific features of the visual scene (e.g. direction, orientation, etc.) (Bae et al., 2018; Goetz et al., 2022). Similarly, in the dLGN, there are at least three broad classes of thalamocortical-projecting (TC) neurons that possess tuning properties similar to those found in RGCs (Krahe et al., 2011; Piscopo et al., 2013). Finally, in V1, there are at least 50-60 cell types that can be discriminated based on genetic, transcriptomic, and physiological markers (Tasic et al., 2018). V1 neurons demonstrate a range of tuning properties, some of which match those found in RGCs and TC neurons, and others that are apparently unique to V1 neurons (Andermann et al., 2011; Wang et al., 2022). Thus, it is well-established that there are neurons with similar tuning properties at each step of the mouse visual pathway.

It is less clear whether neurons tuned to specific visual features are connected to downstream neurons with the same tuning. Indeed, in the mouse, connectivity between the retina and dLGN ranges from highly selective to exceedingly complex (Morgan et al., 2016; Rompani et al., 2017). Nevertheless, there is some evidence that functional segregation exists between the retina and dLGN. TC neurons in the dLGN operate in different ‘modes’, with so-called relay neurons receiving input from a small number of functionally similar RGCs (Rompani et al., 2017). However, the functional response properties of relay-mode TC neurons, as well as the laminar location and tuning properties of the V1 neurons they contact, remain unknown. In addition, direction-selective (DS) and orientation-selective (OS) RGCs project preferentially to the dLGN shell which contains DS and OS TC neurons that deliver tuned input to V1 neurons in superficial layers (Cruz-Martín et al., 2014; Kondo and Ohki, 2016; Marshel et al., 2012). These findings provided evidence for a ‘shell pathway’ dedicated to motion processing, although it is not known whether the V1 neurons contacted by the shell pathway have similar tuning properties. Moreover, there is evidence that DS and OS thalamic input is also delivered to deeper cortical layers, indicating that motion processing occurs outside of the shell pathway. (Sun et al., 2016; Zhuang et al., 2021). Together, these studies provide emerging evidence for functional segregation of tuning properties in retinogeniculate pathways, although the type, laminar location, and tuning properties of the V1 neurons contacted by these pathways remain to be elucidated.

In the primate visual system, there are clearly segregated magno- and parvocellular pathways that carry distinctly different tuning for spatial and temporal frequencies from the retina to different subdivisions of layer 4 (L4) in V1 (Nassi and Callaway, 2009). But the lack of corresponding magno- or parvocellular RGC types and L4 subdivisions in the mouse dLGN, as well as the apparent lack of L4 subdivisions in V1, suggest that parallel pathways for spatiotemporal frequency might not be conserved between mice and primates. On the other hand, the existence of functional segregation for other features in the mouse visual pathway suggests that tuning in V1 could be inherited from downstream neurons, and there is evidence suggesting that this can occur. For example, disruption of direction selectivity in the retina reduces direction selectivity in layer 2/3 (L2/3) V1 neurons, indicating that direction-selective retinal signals contribute to the generation of V1 tuning (Hillier et al., 2017). Moreover, DS and OS TC input is delivered to both superficial and deep layers of V1, suggesting that motion sensitivity in L4 V1 neurons may be influenced by the DS and OS TC inputs they receive (Cruz-Martín et al., 2014; Marshel et al., 2012; Zhuang et al., 2021). At the same time, it has been shown that non-motion-tuned TC input with a specific spatiotemporal offset can create an initial bias for direction preference in L4 V1 neurons (Lien and Scanziani, 2018). Thus, it remains to be determined precisely how motion-tuned TC input is integrated and utilized by V1 neurons. In summary, a great deal is known about how visual information is encoded by cells at each step of the mouse visual pathway. What is currently lacking is a systematic analysis of connectivity that links the tuning properties of the connected neurons at each step of the pathway.

Here, we sought to map the connectivity of L4 V1 neurons with known tuning properties with their presynaptic partners throughout the visual pathway. To this end, we employed two transgenic mouse lines (Nr5a1-Cre and Scnn1a-Tg3-Cre) that label mostly non-overlapping populations of L4 V1 neurons and performed functional, anatomical, and trans-synaptic tracing experiments to characterize their response properties and connectivity. Using 2-photon calcium imaging (2PCI) of GCaMP6s signals to measure responses to drifting sine wave gratings at a range of temporal and spatial frequencies, as well as coherent dot motion stimuli, we found that these L4 V1 neurons differed in their preferred spatial frequency, temporal frequency, and speed tuning, as well their direction and orientation selectivity. Anatomical analyses indicated differences in morphological and laminar position, further establishing that these mouse lines label mostly separate populations of neurons. Lastly, we employed trans-synaptic G-deleted rabies virus (RVdG) tracing to study the RGC types that provide input (via dLGN) to these L4 V1 neurons. We found that each of these L4 V1 neuron types received biased input from different RGC types. Notably, the tuning properties of the traced RGCs matched the tuning of the L4 V1 neurons to which they provide input: More DS V1 neurons received preferential input from DS RGCs (dsRGCs), while more OS V1 neurons received preferential input from OS RGCs (osRGCs).

## Results

### Nr5a1 and Scnn1a neurons prefer different spatial frequencies, temporal frequencies, and speeds

The Nr5a1-Cre and Scnn1a-Tg3-Cre mouse lines have been shown to label restricted populations of L4 V1 neurons (Gouwens et al., 2019; Madisen et al., 2010). We first sought to determine to what extent these mouse lines label functionally distinct populations of V1 neurons by characterizing tuning properties of Cre+ neurons in these mouse lines. To accomplish this, we performed two photon calcium imaging (2PCI) of visual responses of V1 neurons in awake mice (Figure 1A). We drove expression of the calcium indicator GCaMP6s in Cre+ neurons in Nr5a1-Cre (Nr5a1 neurons) and Scnn1a-Tg3-Cre (Scnn1a neurons) mice by injecting AAV-FLEX-GCaMP6s into V1 (Figure 1B). To compare the tuning properties of Nr5a1 versus Scnn1a neurons, we presented drifting sine wave gratings at multiple combinations of SFs and TFs (Figure 1C). For neurons that were visually responsive (see Methods), various response metrics were calculated, including the preferred SF, preferred TF, as well as the preferred TF/SF ratio. The TF/SF ratio was calculated at the preferred SF and TF of each neuron and represents a measure of the neuron’s preferred speed of visual motion (Figure 2). Qualitatively, Scnn1a neurons responded more broadly to a range of SF, TF and TF/SF ratios (Figure 2B). When we compared the preferred responses for both cell types, we found that Nr5a1 neurons preferred lower SFs, higher TFs, and faster TF/SF ratios than Scnn1a neurons (Figure 2C; SF: p = 5.53e-21; TF: p = 7.16e-41; TF/SF: p = 1.52e-48). Thus, our findings provide evidence that Nr5a1 and Scnn1a neurons represent functionally distinct populations of L4 V1 neurons.

**Figure 1:**
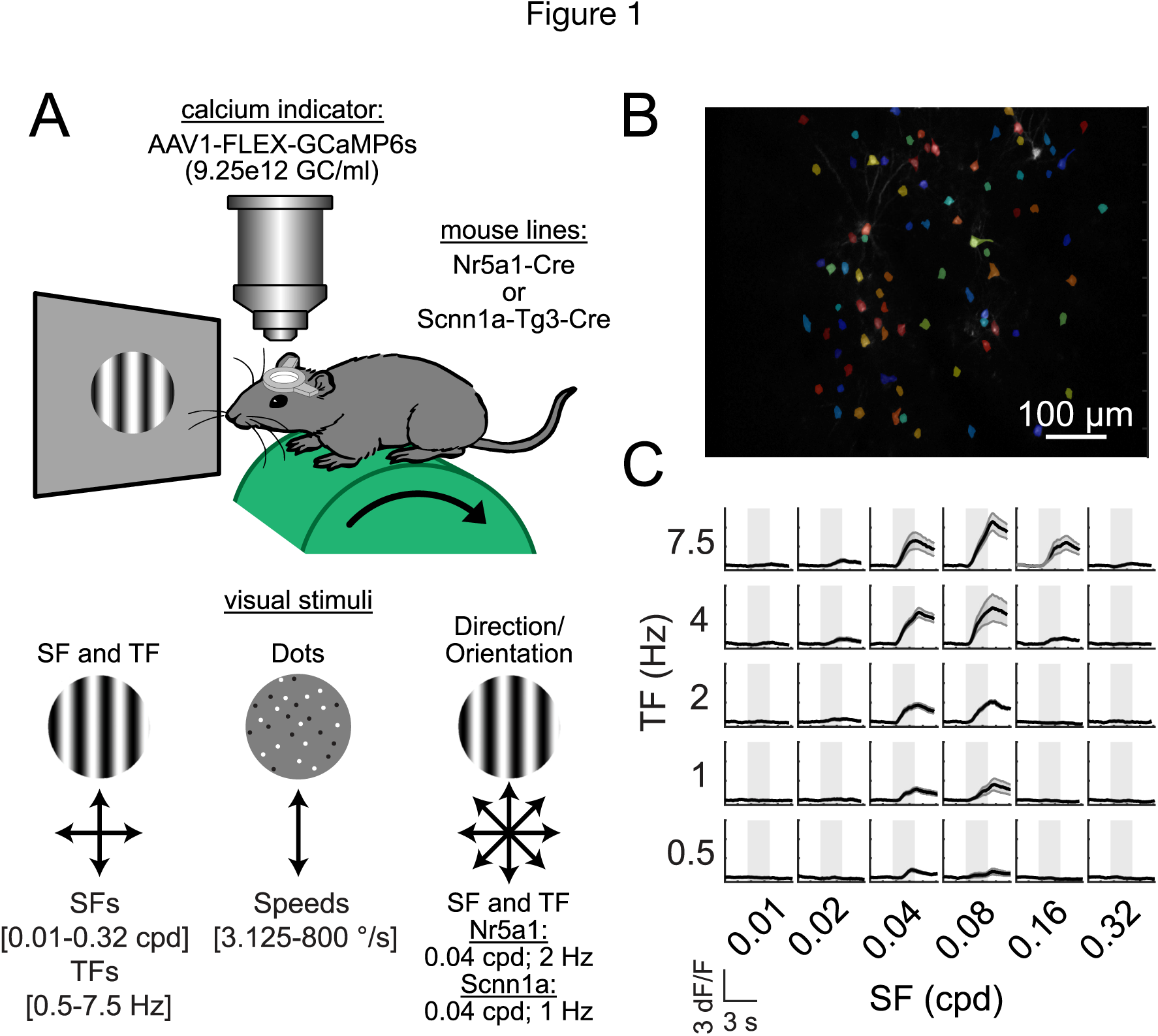
Summary of experimental setup for functional 2PCI and visual stimuli for probing SF, TF, speed, direction, and orientation tuning. (A) Schematic of 2PCI and visual stimulus setup (top) and diagram of stimulus sets used to measure tuning (bottom). Three different stimulus sets were used to measure SF, TF tuning, speed tuning, and orientation/direction tuning. (B) sample field of view in V1, with cell body regions of interests segmented and pseudo-colored randomly. Scale bar = 100 um. (C) Example dF/F responses to different SF and TF combinations presented. Black line represents the mean response, gray shading surrounding the mean corresponds to the standard error of the mean. Shaded gray box corresponds to the 2 second stimulus presentation period.

**Figure 2:**
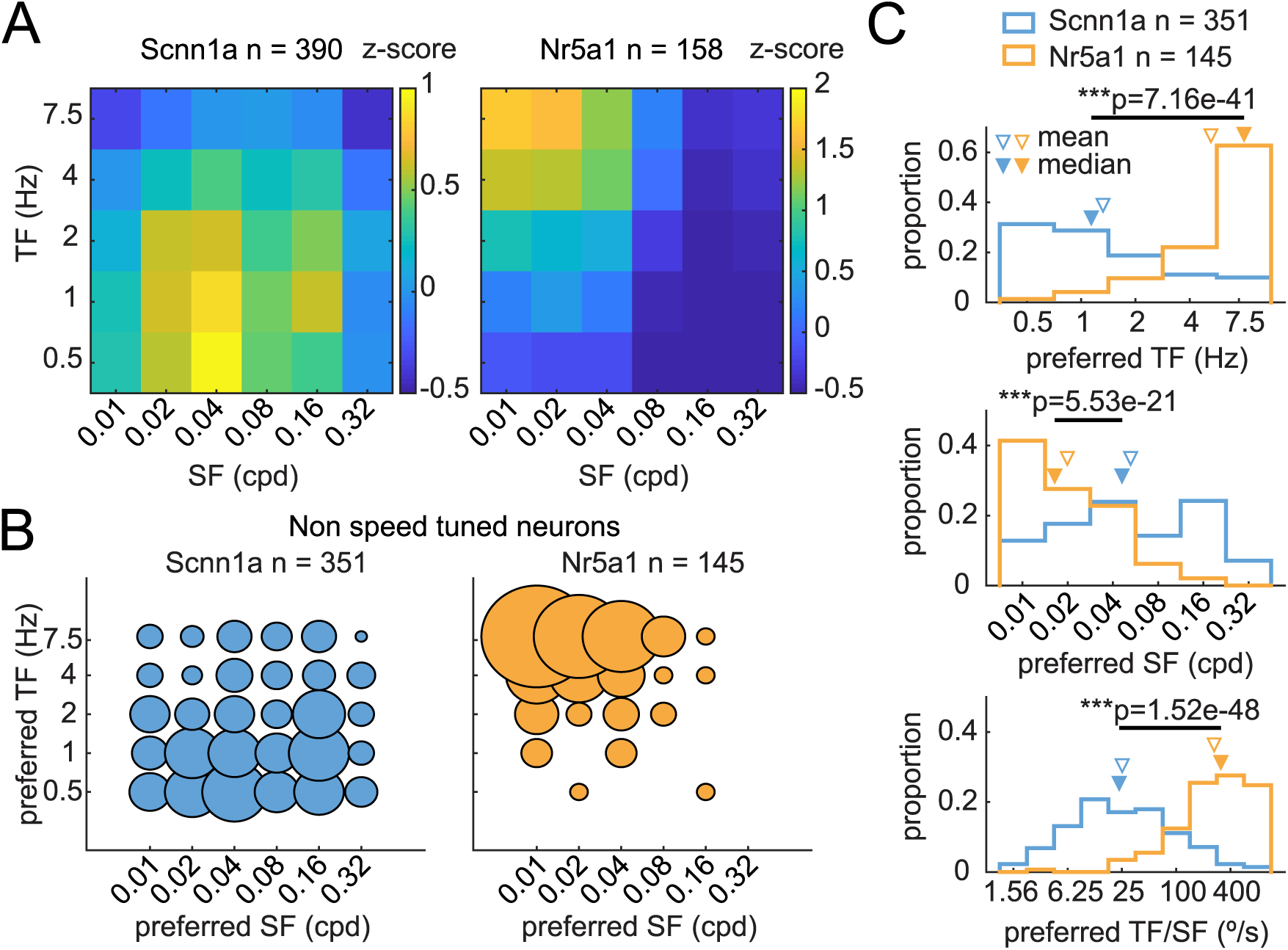
Difference in preferred SF, TF, and speed of visual motion for Nr5a1 and Scnn1a neurons. (A) Mean z-scored responses at preferred orientation to SF and TF combinations presented for all neurons. (B) 2-D Histogram of preferred SF and TF combinations of Nr5a1 and Scnn1a neurons that did not show evidence of speed tuning (see Methods). (C) Histograms of preferred SF, TF, and TF/SF ratio of the same Nr5a1 and Scnn1a neurons as in (B). Nr5a1 neurons preferred lower SFs, higher TFs, and faster speeds (TF/SF ratio). n = total number of cells analyzed. For (C), ***p < 0.001, Wilcoxon Rank-Sum Test.

### Speed tuning of Nr5a1 and Scnn1a neurons

The differences we found in the TF/SF ratios demonstrated that Nr5a1 and Scnn1a neurons preferred different speeds of visual motion. However, these differences do not indicate whether a neuron is speed tuned. The TF/SF ratio is calculated at a single TF and SF, while neurons that are speed tuned prefer a conserved ratio of TF/SF across a range of SFs and TFs (Priebe et al., 2006). When responses of a speed tuned neuron to a range of SFs and TFs are plotted in log_2_ scale, this results in a “ridge” of preferred responses that falls along a line with a slope approximately equal to 1. To assess speed tuning, we fit the responses to a range of SFs and TFs with a modified 2D Gaussian and used parameters of the fit to estimate the speed tuning slope and preferred speed of each neuron (Figure 3A; see Methods). In both mouse lines, there was no difference in the number of speed tuned neurons (∼8-10% of the neurons, see Table 1). However, Scnn1a neurons had significantly higher speed tuning slopes, indicating that they were more speed tuned than Nr5a1 neurons (Figure 3B; p= 0.013). In contrast, we found that Nr5a1 neurons preferred faster speeds than Scnn1a neurons (Figure 3C; p = 7.62e-7). The preferred speeds of Nr5a1 and Scnn1a neurons calculated from the 2D Gaussian fits were in good agreement with the preferred speeds calculated from the TF/SF ratios.

**Figure 3:**
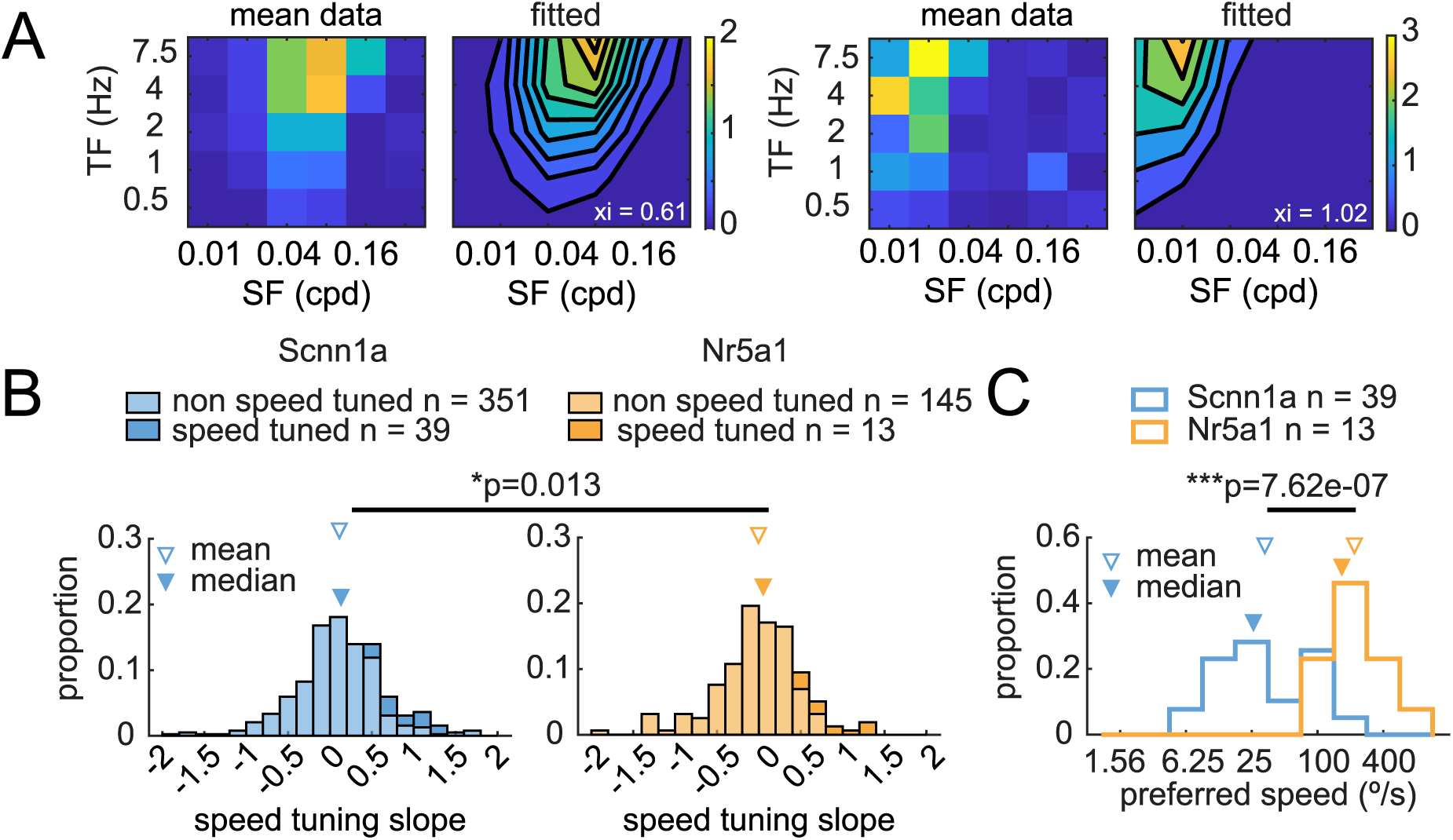
Nr5a1 and Scnn1a neurons possess different speed tuning. (A) Mean dF/F response of two example neurons to a range of SFs and TFs (left) and their fit to the 2-D Gaussian function (right). Fitted speed tuning slope (xi) is displayed in bottom right of each fit. (B) Speed tuning index as measured by slope of 2-D Gaussian fit of Scnn1a versus Nr5a1 neurons. Lighter bars represent non-speed tuned neurons and darker bars represent speed tuned neurons. Scnn1a neurons had stronger speed tuning. (C) Preferred speed of speed tuned Nr5a1 and Scnn1a neurons. Nr5a1 neurons preferred faster speeds. n = total number of cels analyzed. *p < 0.05, ***p < 0.001. Wilcoxon Rank-Sum Test.

**Table 1:** 2-Photon Experiment Numbers Summary. Summary of animals, imaging sessions (Expts), and cells (ROIs) used in calculating tuning preferences.

| Spatial and Temporal Frequency Experiments |  |  |  |  |  |  |  |
| --- | --- | --- | --- | --- | --- | --- | --- |
| Strain | Animals | Expts | ROIs | responsive | % responsive | speed fit | % speed fit |
| Nr5a1 | 5 | 5 | 558 | 285 | 51% | 158 | 55% |
| Scnn1a-Tg3 | 7 | 9 | 4503 | 1426 | 32% | 390 | 27% |
| Coherent Motion Experiments |  |  |  |  |  |  |  |
| Strain | Animals | Expts | ROIs | responsive | % responsive | speed fit | % speed fit |
| Nr5a1 | 4 | 4 | 327 | 54 | 17% | 30 | 56% |
| Scnn1a-Tg3 | 3 | 5 | 1915 | 211 | 11% | 35 | 17% |
| Orientation and Direction Tuning Experiments |  |  |  |  |  |  |  |
| Strain | Animals | Expts | ROIs | responsive | % responsive |  |  |
| Nr5a1 | 4 | 4 | 327 | 158 | 48% |  |  |
| Scnn1a-Tg3 | 3 | 5 | 1915 | 395 | 21% |  |  |

We additionally measured preferred speed using coherent dot motion stimuli and found that, although the overall proportion of neurons responsive to coherent dot motion was small, neurons responsive to, and well-fit by, both dot and sine wave gratings measures of speed tuning were correlated in their preferred speed measured from each stimulus set (Table 1; Supplementary Figure 1A, B, and C; p = 3e-4). Additionally, the tuning preference of preferred speeds was similar to those calculated from sine wave gratings or TF/SF ratios: Nr5a1 neurons preferred faster speeds than Scnn1a neurons based on their fitted speed, as well as upper and lower half max cutoffs (Supplementary Figure 1D and E; fitted speed: p = 5.9e-6; upper half max: p = 2.0e-5; lower half max: p = 2.7e-6). Lastly, we found that Nr5a1 neurons had slightly larger tuning half-widths compared to Scnn1a neurons (Supplementary Figure 1D; p = 0.003). Collectively, our data investigating the speed tuning of Nr5a1 and Scnn1a neurons indicates that a similar portion of both populations of neurons are speed tuned, with Scnn1a neurons having stronger speed tuning and Nr5a1 neurons preferring significantly faster speeds.

### Nr5a1 and Scnn1a neurons possess different direction and orientation selectivities

We also examined the direction and orientation selectivity of Nr5a1 and Scnn1a neurons by performing 2PCI of visual responses to drifting sine wave gratings at 8 different directions. Because the preferred TF of a neuron can depend on the SF of a visual stimulus, we presented the drifting grating stimuli at SF and TF combinations that matched the preferred tuning for each cell type (see Methods) (Mesa et al., 2021; Wang et al., 2022). Both Nr5a1 and Scnn1a neurons demonstrated visual response that were direction- and orientation-selective, and Nr5a1 neurons tended to be more direction-selective than Scnn1a neurons (Figure 4A and B). To quantify the direction and orientation selectivity of these neurons, we calculated the direction selectivity index (DSI) and orientation selectivity index (OSI) for all visually responsive neurons (see Methods). We found that Nr5a1 neurons had significantly higher DSIs than Scnn1a neurons (Figure 4C; p = 1.51e-8), while Scnn1a neurons had significantly higher OSIs than N5ra1 neurons (Figure 4D; p = 5.49e-6). Further, the majority of visually responsive Nr5a1 and Scnn1a neurons were considered direction-selective (DSI > 0.3; Nr5a1: 130/155 neuron; Scnn1a: 203/395 neurons) and orientation-selective (OSI > 0.3; Nr5a1: 151/155 neurons; Scnn1a: 364/395 neurons). Our direction and orientation tuning experiments indicate that Nr5a1 and Scnn1a neurons differ in their tuning to visual motion, with Nr5a1 neurons possessing stronger direction tuning and Scnn1a neurons possessing stronger orientation tuning.

**Figure 4:**
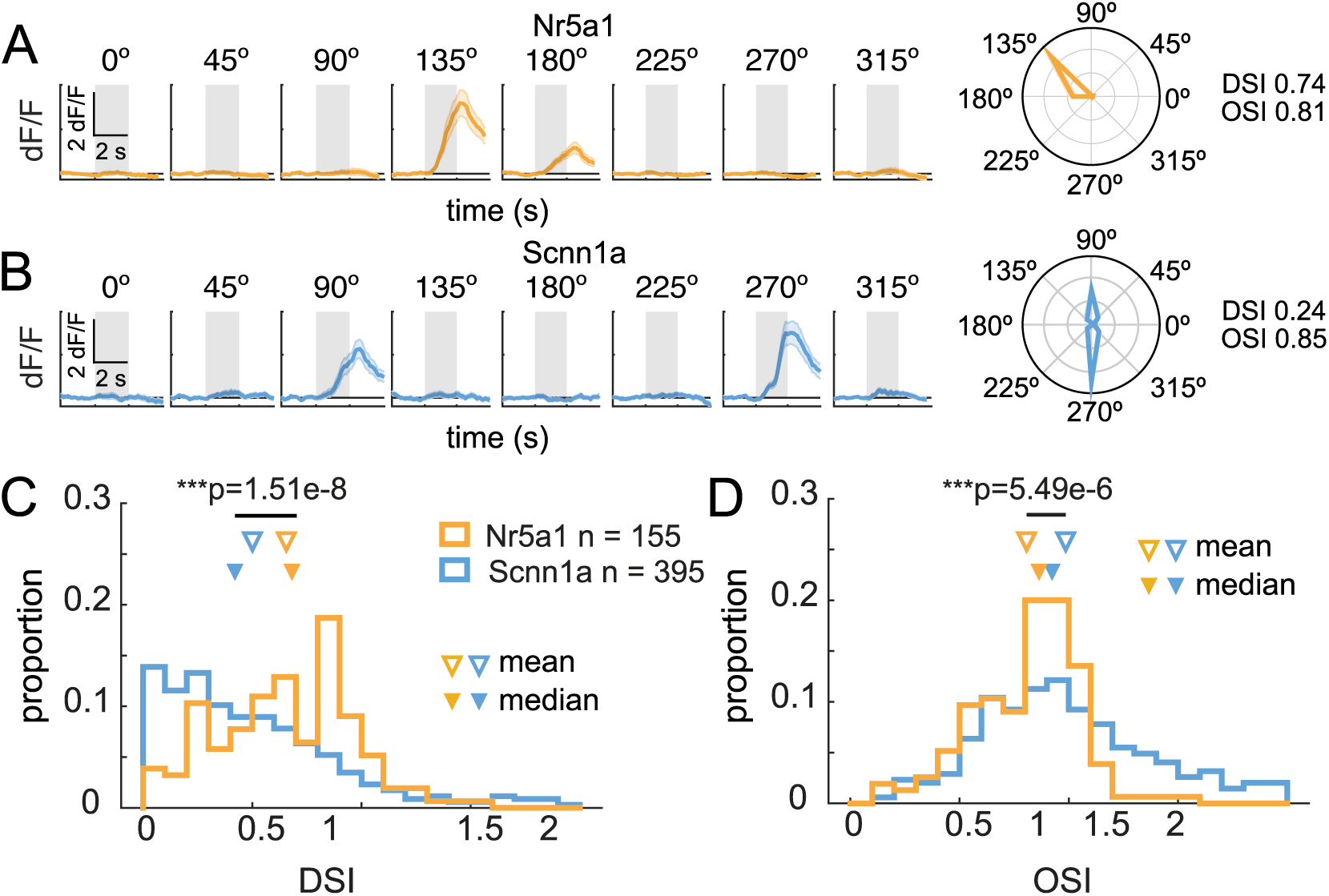
Nr5a1 and Scnn1a neurons differ in the their orientation and direction tuning. (A) Representative dF/F responses of an Nr5a1 neuron to drifting sinusoidal gratings presented at 8 different directions (left), and polar plot summarizing the cell’s tuning with calculated DSIs and OSIs (right). Dark orange line represents the mean response, light orange shading surrounding the mean corresponds to the standard error of the mean. Shaded gray box in dF/F traces corresponds to the 2 second stimulus presentation period. (B) Same as in (A) for a representative Scnn1a neuron. (C) Histogram of proportion of DSIs of Nr5a1 and Scnn1a neurons. (D) Same as in (C) for OSIs. Nr5a1 neurons had higher DSIs, while Scnn1a neurons had higher OSIs. n = total number of cells analyzed and ***p < 0.001, Wilcoxon Rank-Sum Test.

### Differences in retinal input relayed to Nr5a1 and Scnn1a neurons

The differences in tuning properties between Nr5a1 and Scnn1a neurons led us to wonder how these differences might arise. We hypothesized that differences in the type of retinal input these populations of L4 V1 neurons receive from retinogeniculocortical pathways might contribute to the differences we observed in their tuning properties. To determine whether there are differences in how retinal input is delivered to these neurons, we employed Cre-dependent transsynaptic tracing using pseudo-typed RVdG. This method allowed us to characterize the types of RGCs whose input was relayed via the dLGN to Nr5a1 and Scnn1a neurons.

In these experiments AAV8-FLEX-oG and AAV8-FLEX-GFP-TVA were injected into V1, and AAV2-oG-GFP was injected into the dLGN of Nr5a1-Cre and Scnn1a-Tg3-Cre mice. This injection scheme caused Cre+ V1 neurons to express TVA and oG, while cells in the dLGN expressed oG (Figure 5A and B). Three weeks later, EnvA+RVdG-mCherry was injected into V1. This virus could only infect Cre+ V1 neurons expressing TVA. Because these neurons also expressed oG they could assemble functional rabies particles (oG+RVdG) and infect dLGN neurons presynaptic to them (Figure 5B). Since the rabies-infected dLGN neurons also expressed oG and could assemble oG+RVdG, they could infect RGCs presynaptic to them. The end result of this viral tracing design was that RGCs that provided input, via the dLGN, to Nr5a1 and Scnn1a L4 V1 neurons expressed mCherry (Figure 5C).

**Figure 5:**
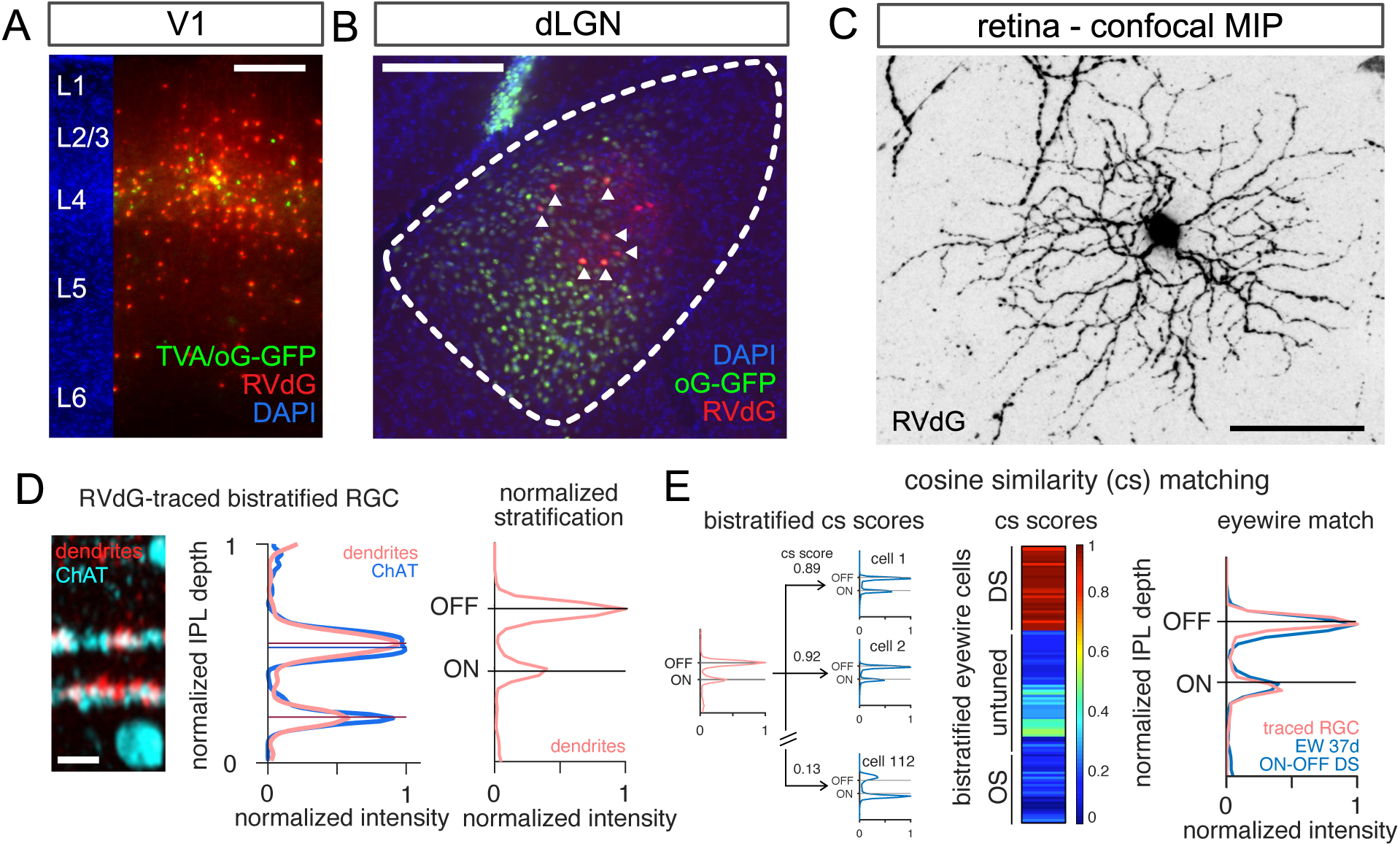
Anatomical reconstruction and cosine similarity matching allows RVdG-traced RGCs to be classified. (A) Coronal sections of V1 showing RVdG (red) and TVA/oG (green) expression in Nr5a1-Cre+ mice. Starter cells are both RVdG+ and TVA/oG+. (B) Coronal section of dLGN showing RVdG (red) and oG (green) expression. Arrowheads indicate RVdG+ cells that also expressed oG-GFP and could retrogradely trace RGCs. (C) Confocal reconstruction of an example RVdG-traced RGC. (D) Z-projection of the stack of confocal images taken from the cell in (C), showing the level of RGC dendrites (red) and ChAT bands (blue, left). The stratification pattern of the cell (middle) can be quantified by normalizing the signal of the dendrites relative to that of the ChAT bands to generate a normalized stratification profile for the cell (right). (E) The RVdG-traced RGC is matched to a specific RGC type by calculating the cosine similarity (cs) between the stratification profile of the RVdG-traced RGC and the stratification profiles of all RGCs in the Eyewire dataset with the same gross stratification pattern (i.e. mono- or bistratified, left). The Eyewire RGC that produces the highest cs score (middle) produces a very good match to the RVdG-traced RGC (right). Scales: 20 µm for retinal cross sections in (D), and 200 µm for all others.

We relied upon the anatomy of the RVdG-traced cells to characterize the types of RGCs that we recovered in these experiments. The stratification position of RGC dendrites within the inner plexiform layer (IPL) has been shown to be a robust way to classify RGC types (Bae et al., 2018; Goetz et al., 2022; Manookin et al., 2008; Sümbül et al., 2014). To determine the stratification position of our RVdG-traced RGCs, the position of the dendrites of each RVdG-traced RGC in the IPL was normalized relative to the choline-acetyltransferase bands (ChAT bands) — two fiduciary bands within the IPL (Figure 5D; see Methods). This allowed us to assign each RVdG-traced RGC to a known RGC type by calculating cosine similarity scores between the normalized stratification profiles of our RVdG-traced cells and the normalized stratification profiles of cells in the Eyewire database (Bae et al., 2018) (Figure 5E; see Methods).

In light of our findings that Nr5a1 and Scnn1a neurons possess different direction and orientation selectivity (Figure 4), we focused our initial analysis on bistratified RGCs. Bistratified RGCs that project to the dLGN are a heterogeneous population of 17 cell types that includes several RGC types tuned to the direction or orientation of motion (Kerschensteiner, 2022) (see Methods). Motion-tuned cells comprise 10 cell types including four types of ON-OFF dsRGCs tuned to the four cardinal directions of motion (dorsal, ventral, nasal, temporal), and six types of osRGCs including three types tuned to horizontal motion and three types tuned to vertical motion, while non-motion-tuned cells comprise the remaining seven bistratified RGC types (Bae et al., 2018; Goetz et al., 2022) (Figure 6A).

**Figure 6:**
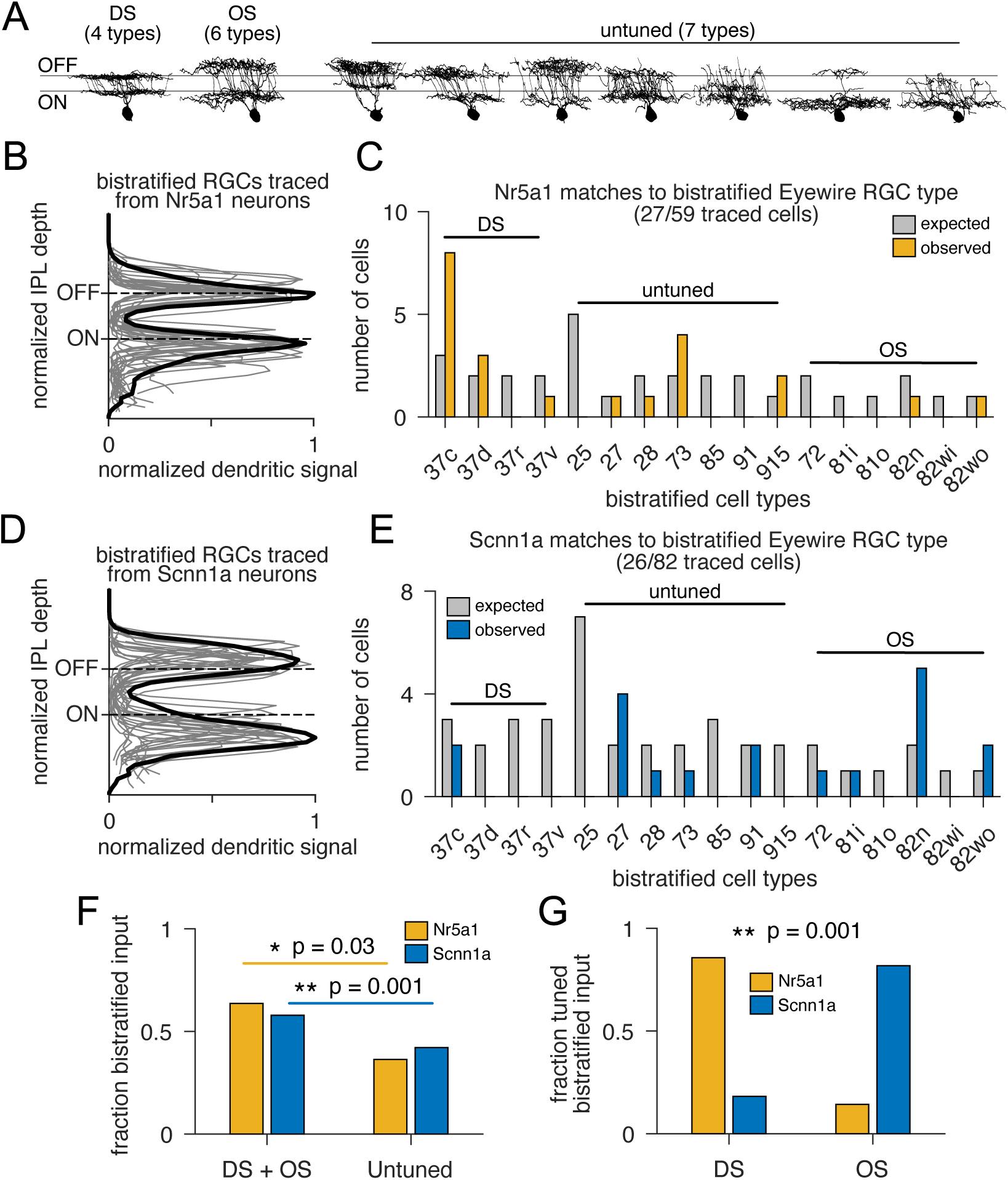
Nr5a1 and Scnn1a neurons differ in the type of input they receive from bistratified RGCs. (A) Example z-projections of the bistratified Eyewire RGC types used for the analysis of bistratified RVdG-traced RGCs (adapted from Bae et al. 2018). (B) Overlays of stratification profiles of individual bistratified RGCs RVdG-traced from Nr5a1 neurons (gray) as well as the normalized mean stratification profile of all cells (black). (C) Distribution of bistratified Eyewire RGC types to which RGCs traced from Nr5a1 neurons were matched. (D and E) Same as in (B and C) for Scnn1a neurons. (F) Fraction of all bistratified input that was DS+OS and untuned for Nr5a1 and Scnn1a neurons. Both Nr5a1 and Scnn1a neurons received more DS+OS than untuned input. (G) Fraction of all tuned input that was DS and OS for Nr5a1 and Scnn1a neurons. Nr5a1 neurons received more DS retinal input, while Scnn1a neurons received more OS retinal input. For (F) Chi Square Test. For (G) Fisher Test. *p<0.05, **p<0.01.

When we analyzed our RVdG-traced RGCs, we found that ∼30-40% of them were bistratified (Nr5a1: 27/59 cells; Scnn1a: 26/82 cells). In addition, the normalized stratification profiles of our bistratified RVdG-traced cells differed between Nr5a1 and Scnn1a neurons, with RGCs traced from Nr5a1 neurons stratifying close to the ON and OFF ChAT bands and RGCs traced from Scnn1a neurons stratifying just below and above the ON and OFF ChAT bands (Figure 6B and D). Further, we found that our bistratified RVdG-traced cells matched to a restricted subset of bistratified RGC types from the Eyewire database for both Nr5a1 and Scnn1a neurons (Figure 6C and E). Collectively, these findings suggested that information from different types of bistratified RGCs is delivered to Nr5a1 and Scnn1a neurons.

To determine whether there were significant differences in the types of bistratified RGCs providing input to Nr5a1 and Scnn1a neurons, we first grouped RGCs into DS and OS (DS+OS) and non-motion-tuned (‘untuned’) categories and determined the fraction of bistratified input that came from each category. We found that both Nr5a1 and Scnn1a neurons received a greater fraction of their bistratified input from DS+OS rather than ‘untuned’ cells (Figure 6F; Nr5a1: p = 0.03; Scnn1a: p = 0.001). Further, there was no difference in the extent of this bias between cell types (p = 0.55). We also examined the fraction of motion-tuned bistratified input that came from dsRGCs or osRGCs. Intriguingly, we found that Nr5a1 neurons were more likely to receive input from dsRGCs, while Scnn1a neurons were more likely to receive input from osRGCs (Figure 6G; p = 0.001). Taken together, these data indicate that Nr5a1 and Scnn1a neurons receive biased retinal input from dsRGCs and osRGCs respectively and, therefore, are relayed different motion-tuned signals from the retina.

We also examined the input that Nr5a1 and Scnn1a neurons received from monostratified RGCs. For our analysis, monostratified RGCs included 28 types of RGCs (13 ON, 11 OFF, 4 ON-OFF) that project to dLGN and do not exhibit robust direction or orientation tuning (Kerschensteiner, 2022) (see Methods). Similar to bistratified RGC input, we found that the majority of our RVdG-traced RGCs matched to a restricted subset of monostratified RGC types (Figure 7A-D). We further quantified this observation by grouping the matched RGCs into three, broad functional categories — ON, OFF, and ON-OFF — and determined the fraction of total monostratified input that came from each of these functional categories. Neither Nr5a1 nor Scnn1a neurons received any monostratified ON-OFF input and, surprisingly, we found that both Nr5a1 and Scnn1a neurons received significantly more input from ON RGCs than OFF RGCs (Figure 7E; Nr5a1: p = 0.007; Scnn1a: p = 2.8e-11).

**Figure 7:**
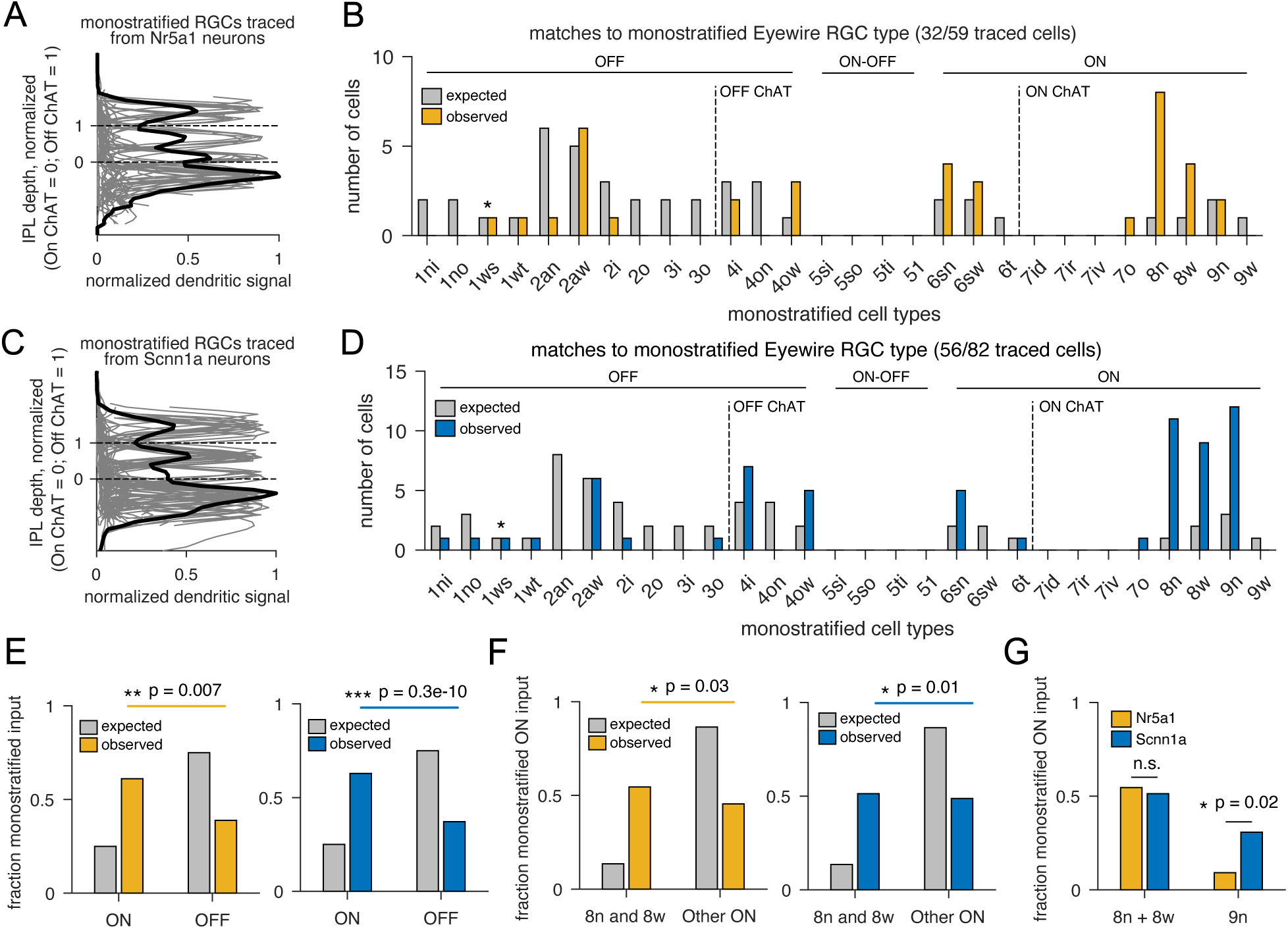
Nr5a1 and Scnn1a neurons receive different patterns of input from ON-type monostratified RGCs. (A) Overlays of stratification profiles of individual monostratified RGCs RVdG-traced from Nr5a1 neurons (gray) as well as the normalized mean stratification profile of all cells (black). (B) Distribution of monostratified Eyewire RGC types to which RGCs traced from Nr5a1 neurons were matched. (C and D) Same as in (A and B) for Scnn1a neurons. Star in (B) and (D) indicates 1ws/M1 RGC that is an OFF-stratifying ON RGC type.(E) Fraction of monostratified input that came from ON and OFF RGCs for Nr5a1 (left) and Scnn1a (right) neurons. Both V1 neurons received more ON than OFF input. (F) Fraction of monostratified ON input to Nr5a1 (left) and Scnn1a (right) neurons that came from Eyewire type 8n and 8w cells versus all other ON RGC types. Both Nr5a1 and Scnn1a neurons received more input from 8n and 8w RGCs. (G) Fraction of monostratified ON input that came from 8n and 8w versus 9n RGCs. Scnn1a neurons received more input from 9n RGCs. For (E) and (G) Chi Square Test. For (F) Fisher Test. *p<0.05, **p<0.01, ***p<0.001.

Of the 13 types of monostratified ON RGCs, the most frequently matched RGC types for both Nr5a1 and Scnn1a neurons were Eyewire types 8w and 8n, with Scnn1a neurons also receiving appreciable input from Eyewire type 9n (Figure 7B and D). Types 8w and 8n likely both correspond to the sustained ON Alpha (sON Alpha) RGC, while type 9n corresponds to the Pix ON RGC (Bae et al., 2018; Goetz et al., 2022) (see Methods). We grouped types 8w and 8n together and compared the frequency with which we recovered them against all the other ON types. Both Nr5a1 and Scnn1a neurons demonstrated a significant bias toward 8w and 8n RGCs (Figure 7F; Nr5a1: p = 0.03; Scnn1a: p = 0.01) and there was no significant difference in the extent of this bias for 8w and 8n cells between cell types (Figure 7G; p = 0.55). In contrast, Scnn1a neurons received significantly more input from 9n RGCs than Nr5a1 neurons (Figure 7G; p = 0.02). While the functional implications of these differences remain to be determined, this is, to our knowledge, the first demonstration of any bias in how input from ON or OFF retinal channels is delivered to V1 neurons in mice.

### Nr5a1 and Scnn1a neurons reside at different depths of layer 4

Lastly, we sought to characterize the proportion of L4 neurons labeled in each mouse line, as well as the depth within V1 at which Nr5a1 and Scnn1a neurons reside. There is evidence in mouse V1 that TC projections with different functional tuning project to slightly different depths of L4, suggesting that V1 neurons residing at different depths of L4 could receive functionally distinct input from dLGN (Zhuang et al., 2021). We crossed Nr5a1 and Scnn1a lines to a Cre-dependent reporter line (Ai14) both alone and as doubly transgenic lines (i.e. Nr5a1-Cre/Scnn1a-Tg3-Cre) and quantified the location of Cre+ V1 neurons. To determine whether these mouse lines only label excitatory neurons, we labeled all neurons (NeuN antibody) as well as inhibitory neurons (GAD67 antibody or mDLX virus, see Methods) and confirmed there was no overlap between Cre+ neurons and inhibitory neurons in either Nr5a1-Cre and Scnn1a-Tg3-Cre mice (Supplementary Figure 2A). Across the depth of V1, Cre+ neurons in all mouse lines peaked between a normalized depth of 0.3-0.5, consistent with labeling in L4 V1 neurons (Supplementary Figure 2B). Interestingly, Nr5a1 neurons were found in more superficial L4, whereas Scnn1a neurons were found deeper in L4 (Nr5a1: 0.38 +/- 0.01; Scnn1a: 0.47 +/- 0.01; p = 0.01; Supplementary Figure 2B). We also calculated the proportion of excitatory L4 neurons labeled by each line and found that Nr5a1 neurons comprised ∼10% of all L4 V1 excitatory neurons, while Scnn1a neurons comprised 49% of all L4 V1 excitatory neurons (Supplementary Figure 2C). In the doubly transgenic Nr5a1/Scnn1a-Tg3-Cre mice, Cre+ neurons labeled 54% of excitatory neurons in L4, suggesting that there is likely a small amount of overlap in the L4 V1 neurons labeled by each mouse line (Supplementary Figure 2C). We next quantified the proportion of Cre+ neurons outside the borders of L4 and found some degree of labeling outside of L4 for each mouse line (Supplementary Figure 2D). Finally, we quantified the apical dendrite length of full dendrite reconstructions of Nr5a1 and Scnn1a neurons in V1 from the Allen Brain Atlas Cell Types database. Scnn1a neurons had longer apical dendrites than Nr5a1 neurons (Supplementary Figure 2E). Taken together, these data indicate that Nr5a1 and Scnn1a represent non-overlapping populations of V1 neurons that possess unique apical morphologies and reside at different depths of L4.

## Discussion

Here, we investigated the retinogeniculocortical connectivity patterns, tuning properties, and morphology of V1 neurons in the Nr5a1-Cre and Scnn1a-Tg3-Cre mouse lines. Collectively, our data indicate that these two mouse lines label different populations of L4 V1 neurons that differ in their tuning properties. Unexpectedly, we found that both of these L4 V1 neuron populations are relayed input from the retina that is biased toward RGCs with tuning properties that matches their tuning. Our main findings are: (1) Nr5a1 neurons are more DS and are relayed more input from dsRGCs. (2) Scnn1a neurons are more OS and are relayed more input from osRGCs. (3) Nr5a1 and Scnn1a neurons receive more input from ON than OFF RGCs, with both receiving input biased toward specific types of ON RGCs. (4) Nr5a1 and Scnn1a neurons differ in their tuning for SF, TF, and speed, with Nr5a1 neurons preferring lower SFs, higher TFs, faster speeds, while being less speed-tuned. (5) Nr5a1 and Scnn1a neurons differ in their cortical depth and apical dendritic morphology, with Nr5a1 neurons residing in a more superficial region of L4 and possessing less extensive apical dendrites (Figure 8). Our findings suggest that specializations exist in the mouse visual system whereby the tuning of neurons at each level of the visual pathway is partly conserved. These specializations likely create an organizational motif that contributes to the generation of different TF, SF, speed, direction, and orientation tuning properties that exist in L4 V1 neurons.

**Figure 8:**
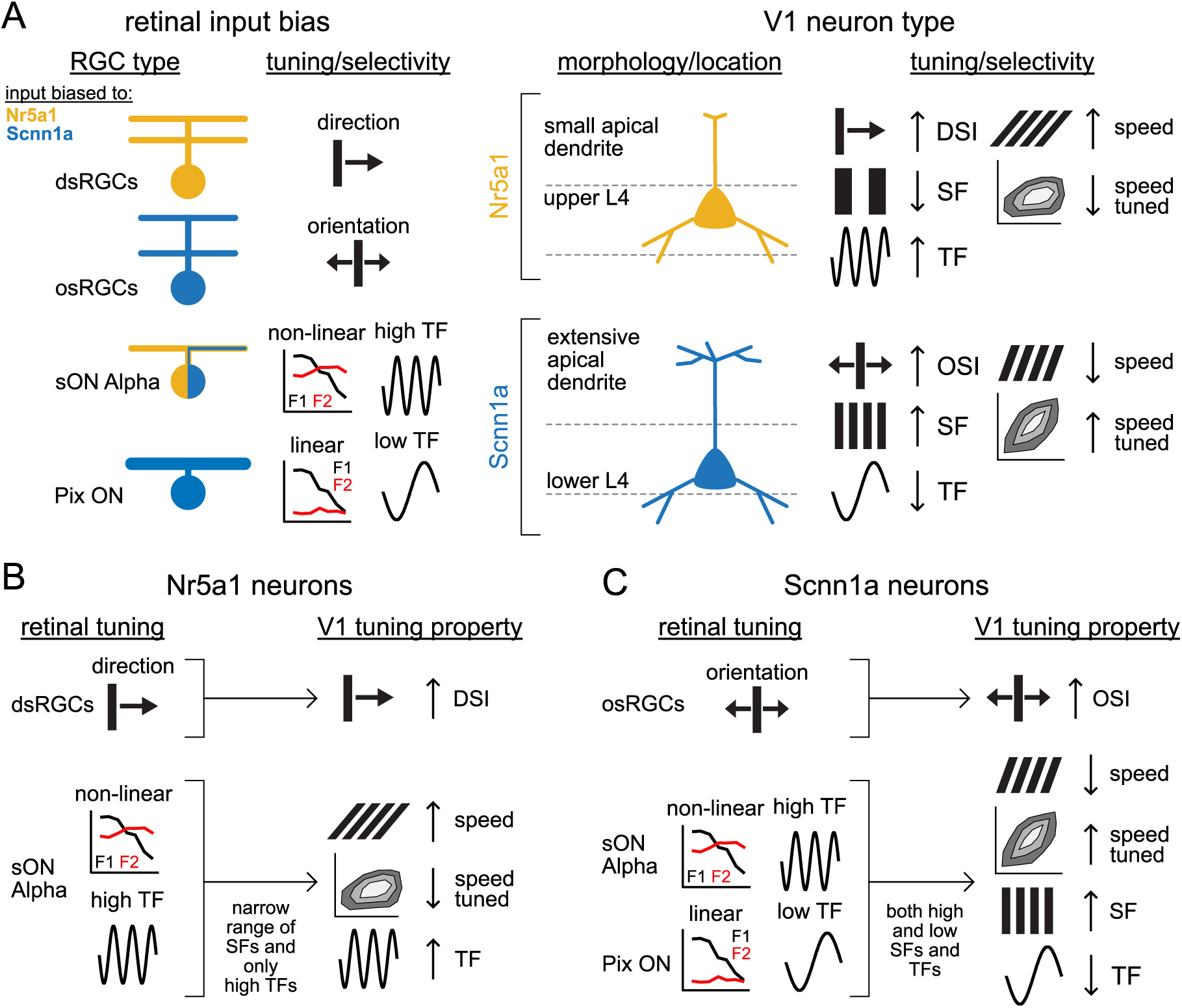
Connectivity, function, anatomy and proposed retinal contribution to tuning in two types of L4 V1 neurons. (A) Nr5a1 and Scnn1a neurons differed in the types of RGCs from which they were relayed biased input (left) as well as their in vivo tuning properties, morphology, and position within L4 (right). Color of RGC in left panel indicates the V1 neuron type to which it sends biased input. (B) Proposed contributions of retinal input to tuning properties in Nr5a1 neurons. Input from dsRGCs likely contributes to strong direction selectivity, while input from sON Alpha RGCs can explain the weaker speed tuning and preference for fast speeds and high TFs seen in Nr5a1 neurons. (C) Same as in (B) for Scnn1a neurons. Input from osRGCs likely contributes to strong orientation selectivity, while the combined input from sON Alpha and Pix ON RGCs can explain the stronger speed tuning as well as the preference for slower speeds, lower TFs, and higher SFs seen in Scnn1a neurons.

### Differences in retinal input relayed to L4 V1 neurons and implications for the generation of tuning in V1

Our findings that the more DS Nr5a1 neurons receive biased input from dsRGCs while the more OS Scnn1a neurons receive biased input from osRGCs represents a significant advancement in our understanding of the organizational motifs within the mouse visual pathway. Recent evidence argued for the existence of two channels in the mouse visual pathway — the core and shell — carrying non-DS and DS information to L4 and L1 V1 layers respectively (Cruz-Martín et al., 2014; Kondo and Ohki, 2016; Marshel et al., 2012). Other studies identified specializations in the dLGN such that TC neurons operate in different ‘modes’, with so-called relay neurons receiving input from a small number of functionally identical RGCs (Rompani et al., 2017). Taken together, these studies suggested that V1 neurons could receive biased retinal input from retinogeniculocortical channels composed of relay neurons carrying different signals routed through the core or shell pathways. However, conflicting reports later emerged showing that superficial and deep layer L4 V1 neurons receive DS and OS inputs from the dLGN (Sun et al., 2016; Zhuang et al., 2021). These findings argued against a clean delineation of the mouse visual pathway into the core and shell pathways and suggested that parallel pathways similar to those in higher mammals are not present in the mouse visual pathway. The data we present here, while agnostic on the precise routing through dLGN, provides additional evidence for the existence of streamlined pathways in the mouse visual pathway carrying DS and OS signals from the retina to V1.

Our findings further suggest that the direction- and orientation-selectivity of V1 neurons could be biased by, or even inherited from, motion-tuned signals originating in the retina. Our data indicate that input from dsRGCs is relayed to strongly DS Nr5a1 neurons, while input from osRGCs is relayed to strongly OS Scnn1a neurons. As outlined above, previous work established that DS and OS TC afferents project to L1 and L4 of V1 (Cruz-Martín et al., 2014; Sun et al., 2016; Zhuang et al., 2021). In the mouse and higher mammals, the response properties of non-motion-tuned dLGN neurons often matches those of the RGCs that innervate them, suggesting retinal input can apparently shape the response properties of dLGN neurons (Suresh et al., 2016; Usrey et al., 1999). Thus, if signals from dsRGCs and osRGCS in the mouse drive DS and OS tuning in the TC afferents they contact, V1 neurons might inherit their direction- and orientation-selectivity directly from the retinal-driven tuning of the TC input they receive. This view is supported by the finding that the elimination of direction selectivity in the retina alters direction selectivity of L2/3 V1 neurons (Hillier et al., 2017). At the same time, because there is evidence that non-motion-tuned TC input can create an initial bias for direction preference in DS L4 V1 neurons, it remains to be seen whether retinothalamocortical inputs to other types of L4 V1 neurons are organized differently (Lien and Scanziani, 2018). Collectively, our data suggest that the mouse visual pathway possesses unique specializations for transmitting DS and OS signals that might contribute to direction- and orientation-selectivity in V1. Notably, this organization differs significantly from that of primates and other higher mammals, where this type of tuning arises de novo in V1 (Priebe, 2016).

### Differences in SF and TF tuning and relation to geniculocortical pathways

The differences we found in the cortical depth as well as the spatial and temporal frequency tuning of Nr5a1 and Scnn1a neurons also support the idea that information delivered by thalamic input is poised to generate tuning properties in V1 neurons. Previous studies demonstrated that afferents targeting superficial parts of L4 were tuned to lower SFs and higher TFs, while afferents targeting deeper regions were tuned to higher SFs and lower TFs (Zhuang et al., 2021). These findings suggested that tuning of L4 V1 neurons can apparently be inherited from the tuning of the thalamic inputs they receive. Consistent with this, we found that Nr5a1 and Scnn1a neurons reside in different L4 sublayers and their tuning properties match those of the TC afferents projecting to those sublayers. Nr5a1 neurons preferred low SFs and high TFs and were found in a more superficial L4 sublayer, while Scnn1a neurons preferred high SFs and low TFs and were localized to a deeper sublayer of L4. Thus, both populations of neurons are positioned in such a way that their SF and TF tuning properties could be generated by the thalamic input they receive. Interestingly, this organizational motif is similar to that seen in primates, with L4 V1 neurons receiving input from the low SF, high TF preferring magnocellular pathway located in the more superficial layer 4Cα, while those receiving input from the high SF, low TF preferring parvocellular pathway located in the deeper layer 4Cβ (Nassi and Callaway, 2009). Collectively, these findings support the idea that the mouse visual pathway possesses some specializations for transmitting information about SF and TF that are similar to those found in higher mammals.

### Differences in speed tuning of L4 V1 neurons and a role for retinal input

Neurons in the mouse visual system are also speed tuned, with neurons in V1 and some higher visual areas (HVAs) tuned to different speeds (Andermann et al., 2011; Wang et al., 2022). Although there is debate as to which HVAs possess stronger speed tuning indices, V1 neurons apparently prefer slower speeds than neurons in HVAs (Wang et al., 2022). However, our analysis of speed tuning in Nr5a1 and Scnn1a neurons suggests that speed tuning in certain L4 V1 neurons can be sharper and faster than previously thought. Existing reports indicated that V1 neurons are tuned to a broad range of speeds centered around 25 °/sec, while neurons in the HVA AL are tuned more sharply to faster speeds around 50-80 °/sec (Andermann et al., 2011; Wang et al., 2022). While Scnn1a neurons preferred speeds that were similar to those previously reported for V1 neurons, we found that Nr5a1 neurons preferred faster speeds that more closely resembled the preferred speed reported for AL neurons. Moreover, Nr5a1 and Scnn1a neurons were easily separated by their preferred speed; a distinction reminiscent of reported differences between neurons in HVAs and V1, rather than between neurons within V1 (Andermann et al., 2011). Thus, our data suggest that tuning for higher speeds emerges earlier in the mouse visual pathway than previously thought. It is worth noting, however, that Nr5a1 neurons represent only ∼10% of all L4 excitatory neurons. Therefore, it remains to be determined whether fast preferred speeds are unique to Nr5a1 neurons, or if other types of L4 V1 neurons exhibit similar speed tuning profiles.

Our findings that Nr5a1 and Scnn1a neurons receive retinal input that is differentially biased toward sON Alpha and Pix ON RGCs provide a potential explanation for the difference in the strength of speed tuning between these neurons. Pix ON and sON Alpha RGC types differ in both their TF tuning and spatial integration properties. Pix ON RGCs integrate information linearly while sON Alpha RGCs possess non-linear spatial summation, and Pix ON RGCs apparently respond to lower TFs than sON Alpha RGCs (Baden et al., 2016; Johnson et al., 2018; Schwartz et al., 2012; Stabio et al., 2018). Cells that integrate spatial information non-linearly can detect small changes in spatial scale within their receptive field — allowing them to resolve higher spatial frequencies — while linear cells cannot (Schwartz et al., 2012). Therefore, a cell receiving input from non-linear and linear cells can detect a broad range of spatial frequencies. Computational models explain the emergence of speed tuning through the convergence of afferents tuned to a range of spatial and temporal frequencies (Simoncelli and Heeger, 1998). Because Scnn1a neurons receive biased input from both sON Alpha (non-linear, higher TFs) and Pix ON (linear, lower TFs) RGCs, their stronger speed tuning could arise through the convergence of retinal input capable of encoding a broad range of both spatial and temporal frequencies. In contrast, weaker speed tuning in Nr5a1 neurons could be the result of convergent retinal input that encodes a narrower range of SFs and TFs. These results, similar to our findings on DS and OS signals, suggest a potential role for retinal input in shaping speed tuning in V1 and warrant experiments probing the contribution of specific RGC types to speed tuning.

## Author Contributions

HW, BKS, and EMC devised the experimental plan. HW conducted experiments and data analysis for imaging and cell counting. BKS and HW conducted rabies-tracing experiments. BKS performed the RGC matching analysis. OD performed cell counting analysis and assisted with perfusions and histology. WN assisted with viral injections, perfusions, and histology. NB assisted with histology and cell counting. HW, BKS, and EMC wrote the manuscript.

## Acknowledgements

This work was supported by the NIH grants EY029932 (HW), T32 GM007198 (HW), EY022577 (EMC), and EY031517 (BKS).

## STAR Methods

### Resource Availability

#### Lead Contact

Further information and requests for resources and reagents should be directed to and will be fulfilled by the Lead Contact, Benjamin Stafford.

#### Materials Availability

This study did not generate new unique reagents.

#### Data and Code Availability

All data reported in this paper will be shared by the lead contact upon request. This paper does not report original code. Any additional information required to reanalyze the data reported in this paper is available from the lead contact upon request.

### Experimental Model And Subject Details

The following transgenic mouse lines were used in this study. For 2PCI experiments, we used Nr5a1-Cre (JAX Stock No: 006364) (n = 4), Ai14/Nr5a1-Cre (n = 1), Scnn1a-Tg3-Cre (JAX Stock No: 009613) (n = 6), and Ai14/Scnn1a-Tg3-Cre (n = 1) transgenic mice ranging in age from P84-115. For quantification of L4 neurons labelled by each line, we crossed Ai14 mice (JAX Stock No: 007914) to generate Ai14/Nr5a1-Cre (n = 3), Ai14/Scnn1a-Tg3-Cre (n = 3), and Ai14/Nr5a1/Scnn1a-Tg3-Cre (n = 3) mice ranging in age from P73-193. For RVdG tracing experiments, we used Nr5a1-Cre (n = 8), Scnn1a-Tg3-Cre (n = 6) transgenic mice ranging in age from P65-85. All experimental procedures followed procedures approved by the Salk Animal Care and Use Committee. Male and female mice were used for all experiments.

### Method Details

#### Surgeries and Viral Injections

For all surgical procedures, mice were anesthetized with either isoflurane (0.5%-2%) or a ketamine/xylazine cocktail (100 mg/kg ketamine, 10 mg/kg xylazine) and secured in a stereotaxic frame. For all surgeries, animals were given analgesics (buprenorphine SR, 0.5-1.0 mg/kg, SQ) at the end of the procedure and provided ibuprofen medicated water (0.11 mg/ml). Intracranial injections were guided by stereotaxic coordinates relative to bregma, anterior-posterior (AP), medial-lateral (ML), and dorsal-ventral (DV).

For 2PCI experiments, mice were implanted with a custom-built circular headframe centered over the left hemisphere. Carprofen (5 mg/kg) and dexamethasone (2 mg/kg) were administered prior to craniotomy surgery. A large 3-5 mm craniotomy was drilled centered over V1 in the left hemisphere and AAV1-Syn-Flex-GCaMP6s-WPRE-SV40 (200-250 nl, 9.25e12 GC/ml, UPenn or 150-200 nl, 7.5e12-1.5e13 GC/ml, Addgene #100845) was injected into the middle of V1 (AP =-3.5 mm, ML=-2.65 mm, DV = -0.35 mm). Following injections, the craniotomy was covered with a glass coverslip (Warner Instruments) mounted on a custom-built ring and sealed to the rest of the headframe with dental cement. Animals were allowed to recover while the virus was expressed over ∼2-3 weeks.

For RVdG tracing experiments, mice were first injected with AAV8-Ef1a-FLEX-GT(Scnn1a: 1.1e11 GC/ml; Nr5a1: 5.4e11 GC/ml; Salk Vector Core; Addgene #26198) and AAV8-CAG-FLEX-oG-WPRE-SV40pA (1.03e13 GC/ml; ViGene; Addgene #74292) mixed 1:2 into the right hemisphere of V1 (AP =-3.4 mm, ML=+2.6 mm, DV=-0.35 mm) at a total volume of 200-250 nl. AAV2-CAG-H2B-GFP-F2A-oG (2.5e11 GC/ml; Salk Vector Core) was injected into the dLGN of the right hemisphere (AP=-2.10 mm, ML =+2.20 DV=-2.45) at a volume of 50-200 nl. 21 days later, EnvA-RvdG-mCherry (3.15e7 TU/mL; Salk Vector Core) at a total volume of 50-200 nl was injected into the previous V1 craniotomy. RVdG was allowed to express 7-10 days before mice were perfused and brains and retinas removed for histology.

To ensure that our RVdG infected starter neurons were specific to the Nr5a1-Cre or Scnn1a-Tg3-Cre labelled V1 neurons, we did controls where we injected only AAV8-FLEX-GT into V1 followed 21 days later by EnvA-RVdG-mCherry virus and confirmed that our starter cell populations were restricted to V1 only (data not shown). Additionally, to ensure that none of the injected AAV2-H2B-GFP-F2A-oG spread retrograde to areas that receive and provide input to dLGN, we also injected AAV2-H2B-GFP-F2A-oG alone into the dLGN of control animals and verified expression was limited to the dLGN (data not shown). In a subset of these experiments, two weeks after injecting AAV2-H2B-GFP-F2A-oG into the dLGN, we also injected VSVg-RVdG-mCherry (150-200 nl; 1.0E7 TU/ml; Salk Vector Core) into V1 to verify there was no retrograde spread of oG back into V1 which would result in transsynaptic spread to dLGN (data not shown).

For a subset of Ai14-crossed mice used in our cell counting analysis, we labelled inhibitory neurons by retro-orbitally injecting AAV-PHP.eB-mdlx-GFP-Fishell-1 into the right eye (50 ul; 6E12 GC/ml; Salk Vector Core; Addgene #83900).

#### In-vivo 2-photon calcium imaging

2PCI was performed on a custom microscope setup which included a Sutter movable objective microscope (Sutter Instruments, Novato, CA) with a resonant scanner (Cambridge Instruments, Bedform, MA). Data acquisition was controlled by a customized version of Scanbox (Neurolabware, Los Angeles, CA). GCaMP6s was excited by a Ti:sapphire laser (Chameleon Ultra II, Coherent, Santa Clara, CA) at 920 nm. Imaging was collected at 1 or 3 planes. For uniplanar experiments, continuous unidirectional scanning was done at 15.49 Hz. For multi-planar experiments, an optotuned lens was used to alternate between depths, and a scanning rate of 5.16 Hz per plane was used. Planes were set ∼20-30 um apart. An area of approximately 500×720 μm (some experiments 800×1230 μm or 570×870 μm) was imaged using a 16x, 0.8 NA objective lens (Nikon Corporation, Tokyo, Japan) through the headframe filled with Immersol-W (Carl Zeiss Microscopy). Prior to imaging, mice were acclimated to the running wheel and visual stimulus setup over 3 days of training sessions.

Running speed was recorded using a rotary encoder. During at least one of these training sessions, GCaMP6s expression was checked in V1. If the imaging field of view (FOV) over the area of expression was obscured due to tissue growth or had poor expression of GCaMP6s, that region was not imaged. For each mouse, we imaged 1 FOV per session, per day.

#### Visual stimulation

Visual stimuli were generated using custom MATLAB code and Psychtoolbox-3 and presented on a gamma corrected monitor (Toshiba 40L5200U, 40’’ or Asus PG279Q, 27’’, both 60 Hz refresh rate) positioned ∼13-23 cm away from the mouse’s right eye. Prior to SF and TF tuning preferences characterization, receptive field locations were mapped by using flashed vertical and horizontal bars, or by manually moving a small drifting stimulus across the monitor, to determine the location that elicited the strongest fluorescent response. The bar receptive field stimuli consisted of vertical or horizontal bars that were 20-21° wide that tiled the entire screen and flashed from black to white over 2 seconds with a TF = 1 Hz and a gray pre/post-stimulus screen and was repeated 10-12 times. Responses to different bar locations were averaged across the entire field of view and the combination of vertical and horizontal bar positions that elicited the strongest response was set as the receptive field center.

SF, TF, and speed tuning were measured using a series of drifting sine wave gratings that varied in 5-6 SFs (0.01-0.32 cycles per degree (cpd) in octave increments), 5 TFs (0.5-7.5 Hz in octave increments), and 2-4 directions (0, 90, 180, 270° or 90, 180°) and were presented in a 40° diameter circular aperture. Each stimulus was repeated 12-15 times. For coherent dot motion experiments, black and white dots 2° in diameter drifted in two directions (90, 180°) within a 40° diameter circular aperture at varying speeds (0, 3.125-800 °/s in octave increments). Dot lifetime was to set 1 second, dot coherence 100%, and dot density 0.2 dots/deg. For orientation and direction tuning experiments, drifting sine wave gratings varied across 8 directions (0-315° in 45° increments) at were presented at a fixed SF and TF that matched the preferred tuning properties for each mouse line (Nr5a1 SF = 0.04 cpd, TF = 2 Hz; Scnn1a SF = 0.04 cpd, TF= 1Hz) within a 40° diameter circular aperture. Each stimulus was repeated 15-20 times. For all experiments, stimulus conditions were presented randomly. For SF, TF, orientation and direction tuning experiments, a 2 second gray screen preceded and followed each 2 second stimulus presentation. For coherent dot motion experiments, the 2 seconds preceding and following the moving stimulus presentation consisted of stationary dots. For all experiments, a stimulus that consisted of a gray screen “blank” was interleaved randomly in 10% of trials. A few experiments (n = 4) had blank trials that consisted of stationary dots during the coherent motion stimulus block, so the “blank” dF/F response was estimated using the pre and post stimulus presentation gray screen.

#### 2-photon calcium imaging processing

Pre-processing was done using suite2p (Version 0.9.0), which included motion correction, cell body region-of-interests (ROI) detection, and neuropil estimation (Pachitariu et al., 2017). ROIs were visually inspected to include only cell bodies and cells that could be well visualized in a max-intensity projection image of the registered frames. The fluorescent trace for each ROI and its corresponding neuropil were then extracted for data analysis. To estimate the contribution of the neuropil to the cell body response, the fluorescent traces were corrected using F_ROI_corrected_= F_ROI_(t)-F_neuropil_(t) (Kerlin et al., 2010). The correction factor, α, was estimated by taking the average ratio of the fluorescence in the blood vessels of the FOV divided by the neuropil. For imaging sessions where 2 or more planes were simultaneously imaged, we identified overlapping ROIs that appeared in multiple planes by calculating the correlation coefficient between the normalized fluorescent signal of all neuron pairs within 25 pixels of each other. ROI pairs with a correlation coefficient greater than 0.4 had the ROI with the lower mean fluorescent signal removed to avoid duplicate ROIs in the dataset.

#### ROI Classification

For each ROI, the response to each trial was calculated by measuring the change in fluorescence from baseline, divided by baseline (dF/F). The baseline for each trial was taken as the mean fluorescence during the 2 second pre-stimulus period. ROIs that were not reliably responsive were eliminated based on 3 criteria. First, the mean fluorescence for the maximum trial condition was less than 6% of the maximum mean fluorescence for ROIs in the FOV. Second, these low fluorescence ROIs had to have a d-prime value (*ẟ* = (*µ_max_* − *µ_blank_*)/(*σ_max_* + *σ_blank_*)) less than 0.5 (Marshel et al., 2011). Finally, some ROIs with extremely high dF/F due to division by a very small baseline were eliminated by removing any ROIs that had a trial dF/F that exceeded the median maximum trial dF/F per ROI + the 95% percentile maximum trial dF/F per ROI. ROIs were determined to be visually responsive for each experiment type by one-way ANOVA with the blank condition included (p<0.05).

#### Quantification of Tuning Responses

All data analysis were analyzed in MATLAB R2018b. For 2PCI experiments, averaged responses to each stimulus condition were calculated by averaging the firing rate or dF/F during the stimulus presentation window. Trials were separated into running versus stationary trials based on the amount of movement that was recorded by the wheel encoder during each trial. For subsequent analysis, only stationary trials (<0.5 cm/s running) were used to avoid any confounds that running may have on neuronal activity (Niell and Stryker, 2010). Overall, approximately 61% of trials were stationary.

#### Estimation of visual tuning properties

SF, TF, and speed tuning was assessed by fitting the trial firing rate or dF/F of each SF and TF combination to a modified 2-dimensional Gaussian function, below using lsqcurvefit in MATLAB (Priebe et al., 2003).

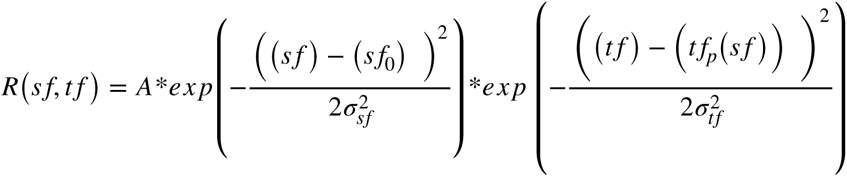

where *t f_p_*_(_*s f* _)_ = *ξ*(*s f* − *s f*_0_) + *t f*_0_

Here A = amplitude of max firing, sf_0_ = preferred SF, tf_o_ = preferred TF, σ_sf_ = SF tuning width (in octaves), σ_tf_ = TF tuning width (in octaves). The tf_p_(sf) variable accounts for the dependence of TF tuning on SF, where ξ is the speed tuning index/slope. When ξ = 1, the neuron is perfectly speed tuned as SF and TF vary in proportion. When ξ = 0, the cell is considered not speed tuned, as SF and TF tuning occur independently, and the preferred speed depends on the SF and TF. For neurons with high directional selectivity (DSI>0.4), only SF and TF trials in the preferred direction are included. Similarly, for neurons with orientation selectivity (OSI>0.2), only SF and TF trials at the preferred orientation were included. Trial responses below zero were rectified to zero prior to fitting. For neurons responsive to all directions, all SF and TF trials were included for model fitting. 95% confidence intervals (CI) were generated by sampling with replacement 500 times. Only neurons that could be well-fit by the 2-D Gaussian were included for subsequent analysis. This criterion required that the 95% CI be less than 3 octaves for the SF_0_, TF_0_, and σ_SF_, σ_TF_ eliminating approximately 40-70% of ROIs and single-units that were not well-fit (see Table 1 and Table 2). Neurons were considered ‘speed tuned’ if the 95% CI for the speed tuning slope included a slope of 1, but not 0. All other cells were classified as ‘not speed tuned’. The median value from the fits was taken as each neuron’s speed tuning index and preferred speed (tf_0_/sf_0_). Sample fits are shown in Figure 3A.

Responses to coherent dot motion were estimated by fitting responses to each presented speed at the preferred direction to a smoothing spline in MATLAB with the smoothing parameter set at 0.3. Neurons were considered well-fit if the 95% confidence interval (CI) for the preferred speed was less than 3 octaves. CIs were generated by sampling with replacement 500 times. From the fits, the preferred speed, upper and lower half max, and tuning half-widths were calculated. For neurons that were high or low pass, the upper and lower half max were set to NaN. A sample spline fit is shown in Supplementary Figure 1A.

Orientation selectivity index and direction selectivity index (OSI and DSI) were calculated in the same manner as previous papers (Marshel et al., 2011).

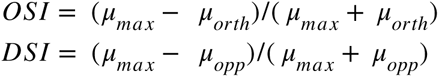

where μ_max_ is the response at the preferred orientation/direction, μ_orth_ is the averaged response of the 2 directions orthogonal to the preferred, and μ_opp_ is the response opposite to the preferred. Because data was collected across multiple imaging sessions, we calculated tuning parameters for each imaging session separately (Supplementary Figure 3). This analysis demonstrated that differences in tuning properties between Nr5a1 and Scnn1a neurons were consistent across imaging days and sessions.

#### Histology

Animals were euthanized by intraperitoneal injection of euthasol (>100 mg/kg) and then perfused with phosphate-buffer saline (PBS), followed by 4% paraformaldehyde (PFA). Brains were removed from skulls and post-fixed in 2% PFA and 15% sucrose at 4C for 24 hours before being transferred to 30% sucrose at 4C for at least another 24 hours. Eyes were removed and placed directly in 4% PFA at 4°C overnight. Retinas were then dissected out and transferred to 1X PBS prior to immunostaining. Brains were sectioned coronally into 50 µM thick sections and then subsequently immunostained followed by DAPI staining. Brain sections were incubated with primary antibody at 4° overnight in 1% normal donkey serum/0.1% Triton-X 100 in PBS, followed by secondary antibody for 2 hours at room temperature. Retinas were blocked over night at 4°C in 10% donkey serum/0.1% Triton-X 100, then placed in primary antibody in the same blocking solution and rocked at 4°C for 72 hours. After washing in 1X PBS, retinas were incubated in secondary antibody diluted in 1X PBS overnight, followed by washing in 1X PBS. Retinas were mounted and cover slipped in Vectashield + DAPI (H-1200-10, Vector Labs).

For 2PCI brains, immunohistochemistry was performed to verify GCaMP6s viral expression by incubating sections with chicken anti-GFP (1:1000, GFP-1020, Aves Labs) followed by Alexa 488 donkey anti-chicken secondary (1:1000-1:500, 703-545-155, Jackson ImmunoResearch).

For cell counting sections, antigen retrieval was performed using 10 nM sodium citrate buffer and then sections were immunostained with rabbit anti-NeuN (1:1000, ab177487, Abcam), mouse anti-GAD67 (1:1000, MAB5406, MilliporeSigma), rat anti-RFP (1:1000, 5F8-100, Chromotek) followed by Alexa 488 donkey anti-rabbit (1:1000, A21207, Thermo Fisher), Alexa 647 donkey anti-mouse (1:500, A31571, Thermo Fisher), and CF568 donkey anti-rat (1:500, 89138-546, Biotium), secondaries. For cell counting brains where inhibitory neurons were labelled with mDlx virus, antigen retrieval was performed as before and then sections were immunostained with rabbit anti-NeuN (1:1000, ab177487, Abcam), chicken anti-GFP (1:1000, GFP-1020, Aves Labs), and rat anti-RFP (1:1000, 5F8-100, Chromotek) followed by Alexa 647 donkey anti-Rabbit (1:1000, A31573, Thermo Fisher), Alexa 488 donkey anti-chicken secondary (1:1000, 703-545-155, Jackson ImmunoResearch), and CF568 donkey anti-rat (1:500, 89138-546, Biotium) secondaries. Sections were first imaged on an Olympus BX63 microscope using a 10x/0.4 NA objective (Olympus) to identify the borders of V1 and layer 4. Sections for cell counting were then imaged on a Zeiss LSM880 confocal microscope using a 20x/0.8NA objective. A stitched z-stack image was taken so that the entirety of V1 in one of the hemispheres of that section could be used for quantification.

For RVdG-tracing experiments, coronal brain sections were immunostained with rabbit anti-ds Red (1:1000, 632496, Takara) goat anti-GFP (1:500, RL600-101-215, Rockland) followed by Alexa 594 donkey anti-rabbit (1:1000, A21207, Thermo Fisher) and Alexa 488 donkey anti-goat (1:1000, A11055, Thermo Fisher) secondaries. Some sections were additionally stained with mouse anti-PV (1:1000, P3088, Sigma-Aldrich) followed by Alexa 647 donkey anti-mouse (1:1000, A31571, Sigma-Aldrich) secondary to visualize the TRN and label inhibitory neurons in the dLGN.

RVdG-traced brain sections were imaged on an Olympus BX63 microscope using a 10x/0.4 NA objective to verify trans synaptic spread of RVdG from V1 to dLGN. For sections with dense labeling, z-stack images were taken and a max-intensity projection image was used to assess viral expression. We additionally examined areas where volume spread of AAV2-H2B-GFP-oG coupled with trans-synaptic RVdG spread from dLGN could result in neurons co-expressing oG and RVdG that could confound our study (eg. TRN, vLGN, and IGL). Any animals in which co-expression of oG and RVdG was seen in retinorecipient structures outside the dLGN were excluded from further analysis.

Retinas from RVdG-tracing experiments were immunostained with rabbit anti-dsRed (1:1000, 632496, Takara) and goat anti-ChAT (1:100, AB144P, Millipore) followed by Alexa 594 donkey anti-rabbit (1:1000, A21207, Thermo Fisher) and Alexa 488 donkey anti-goat (1:1000, A11055, Thermo Fisher) secondaries. Labelled RGCs were imaged on a laser scanning confocal (Zeiss LSM710) using a 40x water immersion objective. Z-stacks were acquired through the full depth of the inner plexiform layer.

#### Cell Counting Quantification

We used ImageJ and the Cell Counter plugin to quantify the number of neurons in L4 of V1 in each imaged section. We quantified the number of L4 neurons (NeuN), inhibitory neurons (GAD67 stain or mDlx virus), and tdT+ neurons to estimate the proportion and depth of L4 excitatory neurons labelled in Nr5a1-Cre, Scnn1a-Tg3-Cre, and Nr5a1/ Scnn1a-Tg3-Cre mouse lines. We additionally counted all tdT+ neurons across the entire depth of V1 cortex since both mouse lines contain a small amount of non-L4 expression. Counters were blind to the strain of the mouse of the sections they were counting. We found that both the GAD67 antibody and mDLX viruses labelled similar proportions of L4 neurons, so we analyzed these sections together. Approximately 1-3 sections containing V1 from each mouse were quantified.

To estimate the depth of Nr5a1 and Scnn1a neurons within V1, we used ImageJ to first draw a line approximating the top and bottom edge of V1 in each section. Next, we created a Euclidean distance map based on these borders. We then used the Euclidean distance maps and the coordinates of each marked cell from the Cell Counter to calculate the distance of each tdT+ cell from the top and the bottom of the cortex. To account for variations in cortical thickness across sections, we normalized the depth as a fraction of the total thickness of cortex at that point (depth from top/ (depth from top + depth from bottom).

#### Rabies Virus Tracing Quantification

We characterized the RGCs labeled from RVdG-tracing experiments by quantifying the stratification position of each cell’s dendrites relative to the choline acetyltransferase bands (‘ChAT bands’) in the IPL as described previously (Manookin et al., 2008; Manookin et al., 2010). Z-stack images of the dendrites of each RGC were acquired in distal regions of the ganglion cell dendritic tree at which point the dendrites were well stratified to minimize the effect of tissue warping. Maximum intensity projections of the dendrites (594 signal) and ChAT bands (488 signal) were created and loaded into custom programs written in Matlab for semi-automatic analysis. To identify the boundaries of the IPL, user-selected points were used corresponding to the x–y position just scleral to the ChAT cell bodies on the vitreal side of the IPL, and just vitreal to the ChAT cell bodies on the scleral side of the IPL. Using the first selected set of x-y coordinates, a line scan was performed along the y-axis at 10 consecutive positions (pixels) along the x-axis in both channels and averaged. This generated profiles of the fluorescence signals of both the dendrites and ChAT bands in the same region of the IPL. The position of the dendrites in the IPL was normalized relative to the ChAT bands such that the ON ChAT band = 0, and the OFF ChAT band = 1. This procedure generated a normalized stratification profile for each cell that allowed comparisons to be made across many cells from different animals. Traced cells were classified as monostratified or bistratified based on whether the normalized stratification profile had one or two peaks.

To match each RVdG-traced cell to a known RGC type, the normalized stratification profile was compared against those from the Eyewire dataset (Bae et al., 2018). Stratification profiles of Eyewire cells were first normalized in the same way as the RVdG-traced cells (ON ChAT = 0; OFF ChAT = 1), and cosine similarity scores were calculated between an RVdG-traced RGC and every cell from the dataset with the same classification (i.e. monostratified or bistratified). The cell from the dataset that produced the highest cosine similarity score was the RGC type to which we matched our traced cell. For the matching process, bistratified RGCs comprised 17 total types which included the following Eyewire types: 25, 27, 28, 37c, 37d, 37r, 37v, 72, 73, 81i, 81o, 82n, 82wi, 82wo, 85, 91, and 915. Monostratified RGCs comprised 28 total types which included the following Eyewire types: 1ni, 1no, 1ws, 1wt, 2an, 2aw, 2i, 2o, 3i, 3o, 4i, 4on, 4ow, 5si, 5so, 5ti, 51, 6sn, 6sw, 6t, 7id, 7ir, 7iv, 7o, 8n, 8w, 9n, 9w.

#### Matched RGC Analysis

To quantify the types of RGCs that were traced from Nr5a1 and Scnn1a neurons, we compared different subsets of RGCs from the Eyewire database. These subsets included the bistratified and monostratified RGCs outlined above. We also compared the number of dsRGCs and osRGCs that were traced. The dsRGCs comprised the following types: 37c, 37d, 37r, 37v, while the osRGCs comprised the following types: 72, 81i, 81o, 82n, 82wi, 82wo. To determine which Eyewire types corresponded to osRGCs, we relied on matches of RGC types from the RGC Typology Project to the Eyewire dataset (Goetz et al., 2022). For statistical testing of matched RGCs, we used the total number of cells in the Eyewire dataset to generate the expected frequency for each RGC type. RGC types that do not project to the dLGN were excluded when calculating expected frequencies (Kerschensteiner, 2022) (Supplementary Figure 4). Expected numbers of RGCs were calculated separately for Nr5a1 and Scnn1a neurons and scaled to the total number of RVdG-traced cells recovered from each mouse line.

For our quantification of monostratified ON RGCs, we considered both Eyewire types 8n and 8w to represent sustained On Alpha (sON Alpha) RGCs. Bae, et al. classified type 8w as ‘securely known’ as the canonical sON Alpha, while type 8n was unmatched. However, the primary distinction between the types was soma size with the authors suggesting they ‘are nearly identical…in stratification and visual response.’ (Bae et al., 2018) Further, the RGC Typology Project failed to find a functional and morphological match to 8n (Goetz et al., 2022). Given the similarity in morphology, stratification, and function between Eyewire types 8w and 8n, it seems reasonable to assume they correspond to the same cell type.

#### Estimation of Apical Dendrite Length

Cell morphology reconstructions from Nr5a1 and Scnn1a neurons in V1 were downloaded from the Allen Brain Atlas Cell Types database (Dataset: Allen Institute for Brain Science, 2015; Gouwens et al., 2019). Only cells where the apical dendrite was considered intact were used. Apical dendritic lengths were measured using the simple neurite tracer plugin in ImageJ.

#### Experimental Design and Statistical Analysis

The values of n and what n represents were reported in results and figure legends. For 2PCI, group differences were determined to be statistically significant by Wilcoxon Rank-Sum Test using MATLAB. For the correlation between preferred speed by coherent dots versus gratings, significance was determined by an F-test comparing the linear model versus a constant model (MATLAB fitlm function). For RVdG-traced RGC analysis, statistical significance was determined by Chi-Square Tests when minimum expected values exceeded 5 cells using MATLAB, otherwise Fisher Exact Tests were used. Comparisons were significant if p < 0.05.

---

| REAGENT or RESOURCE | SOURCE | IDENTIFIER |
| --- | --- | --- |
| Antibodies |  |  |
| Chicken anti-GFP | Aves Labs | Cat # GFP-1020 |
| Mouse anti-PV | Sigma Aldrich | Cat # P3088 |
| Rabbit anti-NeuN | Abcam | Cat # ab177487 |
| Mouse anti-GAD67 | Millipore Sigma | Cat # MAB5406 |
| Rat anti-RFP | Chromotek | Cat # 5F8-100 |
| Goat anti-GFP | Rockland | Cat # RL600-101-215 |
| Rabbit anti-dsRed | Takara | Cat # 632496 |
| Goat anti-ChAT | Millipore Sigma | Cat # AB144P |
| Donkey anti-Rabbit, Alexa Fluor 647 | ThermoFisher | Cat # A31573 |
| Donkey anti-Rat, CF 568 | Biotium | Cat # 89138-546 |
| Donkey anti-Mouse, Alexa Fluor 647 | ThermoFisher | Cat # A31571 |
| Donkey anti-Rabbit, Alexa Fluor 594 | ThermoFisher | Cat # A21207 |
| Donkey anti-Goat, Alexa Fluor 488 | ThermoFisher | Cat # A11055 |
| Donkey anti-Rabbit, Alexa Fluor 488 | ThermoFisher | Cat # A21207 |
| Goat anti-ChAT | Millipore Sigma | Cat # AB144P |
| Donkey anti-Chicken, Alexa Fluor 488 | Jackson ImmunoResearch | Cat # 703-545-155 |
| Bacterial and virus strains |  |  |
| AAV1-Syn-Flex-GCaMP6s-WPRE-SV40 | UNC Vector Core/Addgene | Addgene # 100845 |
| AAV8-Ef1a-FLEX-GT | Salk Vector Core GT3 | Addgene # 26198 |
| AAV8-CAG-FLEX-oG-WPRE-SV40pA | ViGene | Addgene # 74292 |
| AAV2-CAG-H2B-GFP-F2A-oG | Salk Vector Core GT3 | N/A |
| EnvA-RvdG-mCherry | Salk Vector Core GT3 | N/A |
| VSVg-RVdG-mCherry | Salk Vector Core GT3 | N/A |
| AAV-PHP.eB-mdlx-GFP-Fishell-1 | Salk Vector Core GT3 | Addgene # 83900 |
| Chemicals, peptides, and recombinant proteins |  |  |
| 4 or 5 mm Glass cover slips | Warner Instruments | Cat # CS-4R/5R |
| Dental Cement | Caulk Dentsply | Cat # 675571 |
| DAPI | ThermoFisher | Cat # D1306 |
| Triton X-100 | ThermoFisher | Cat # AC327371000 |
| Vectashield | Vector Labs | Cat # H-1200-10 |
| Experimental models: Organisms |  |  |
| Mouse: Nr5a1-Cre | Gift from Dr. Hongkui Zheng, Allen Institute | JAX Stock # 006364 |
| Mouse: Scnn1a-Tg3-Cre | Jackson Labs | JAX Stock # 009613 |
| Mouse: Ai14 | Jackson Labs | B6.Cg-Gt(ROSA)<br>26Sor<sup>tm14</sup>(CAG-tdTomato)<sup>Hze</sup>/J<br>JAX Stock # 007914 |
| Software and algorithms |  |  |
| MATLAB 2016/2018b | Mathworks | https://www.mathworks.com/products/matlab.html |
| Zen | Zeiss | N/A |
| CellSens | Olympus | N/A |
| FIJI | NIH ImageJ Software | https://fiji.sc |
| ImageJ | NIH ImageJ Software | https://imagej.net/ij/index.html |
| Scanbox | Neurolabware | http://scanbox.org |
| Suite2p | Pachitariu, et al., 2017 | https://www.suite2p.org/ |
| Other |  |  |
| BX63 microscope | Olympus | N/A |
| LSM710 microscope | Zeiss | N/A |
| Airyscan 880 microscope | Zeiss | N/A |
| Movable objective microscope | Sutter Instrumenst | N/A |
| Chameluon Ultra II Ti-sapphire laser | Coherent | N/A |
| Resonant scanner | Cambridge Instruments | N/A |
| Sliding Microtome | Microm | N/A |

