## Supplemental Figures for "Parallel pathways carrying direction and orientation selective retinal signals to layer 4 of mouse visual cortex"

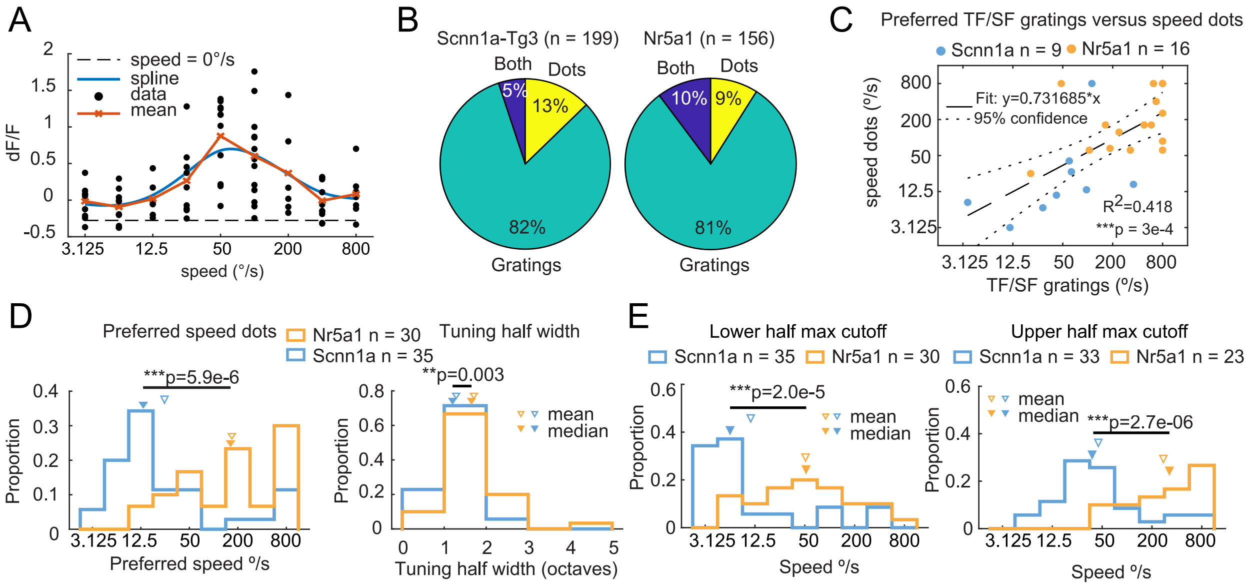


Supplementary Figure 1: Coherent dot motion speed tuning preferences. (A) Example neuron response to dot motion and spline fit. Black dots are the individual trial dF/F responses, red line is the mean response per combination of speed, dotted black line is the mean response to speed = 0 °/s, and blue line is the spline fit. (B) Proportion of neurons well-fit by speed tuning versus coherent motion speed or both.(C) Comparison of preferred speed as measured by sine wave gratings versus coherent dots for neurons fit and responsive to both stimuli. Linear model fit with intercept. (D) Preferred speed (left) and tuning width (right) of all neurons responsive and well-fit to coherent motion stimulus. Nr5a1 neurons preferred faster speeds. (E) Lower and upper half-max cut offs for preferred speed by coherent motion stimulus. For (B-D) n = total number of cells analyzed. **p < 0.01, ***p < 0.001. For (C) F-test. For (D) and (E) Wilcoxon Rank-Sum Test.

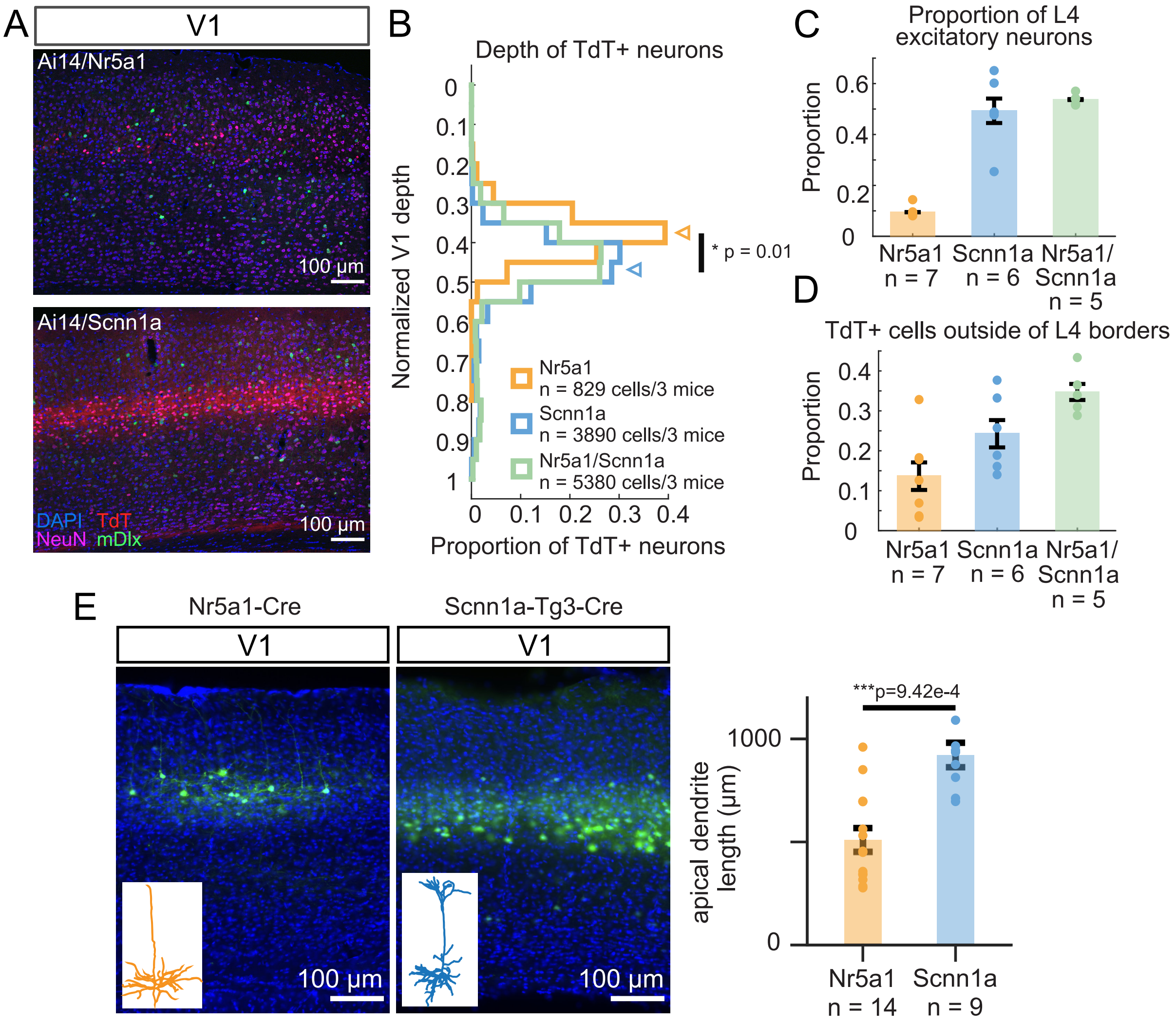


Supplementary Figure 2: Quantification of proportion of layer 4 neurons labelled by Nr5a1 and Scnn1a mouse lines. (A) Sample section from z-stack images used for cell counting. Scale bar = 100 µm. (B) Normalized depth of TdT+ neurons in Ai14/Nr5a1, Ai14/Scnn1a, and Ai14/Nr5a1/Scnn1a mice. (C) Proportion of excitatory neurons labelled by each Ai14 cross in layer 4. (D) Proportion of TdT+ neurons in each mouse line found outside of layer 4 borders. For (C) and (D), n = number of sections quantified, and error bars represent standard error of the mean. (E) AAV1-FLEX-GCaMP6s expression in Nr5a1-Cre (left) and Scnn1a-Tg3-Cre (middle) mice in V1, and quantification of apical dendritic length (right) of Nr5a1 and Scnn1a neurons. Inset images in (E) are sample dendritic reconstruction of a L4 cell from each mouse. Data in (E) came from Allen Brain Atlas of Cell Types. For (B) reported p-value is from comparison between animals. (E) n = total number of cells analyzed and error bars represent standard error of the mean between animals. For (C) and (D) n = number of sections analyzed. $^*p < 0.05$, $^{***}p<0.001$, t-test.

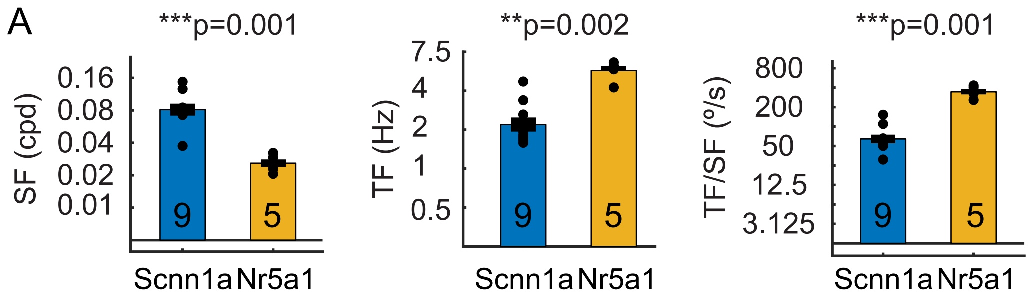


Supplementary Figure 3: Tuning properties of Nr5a1 and Scnn1a neurons are stable across multiple imaging sessions. Mean preferred SF (left), TF (middle), and TF/SF ratio (right) per imaging session. Error bar denotes standard error of the mean (SEM).**$p<0.01$, ***$p<0.001$, Wilcoxon Rank-Sum Test.

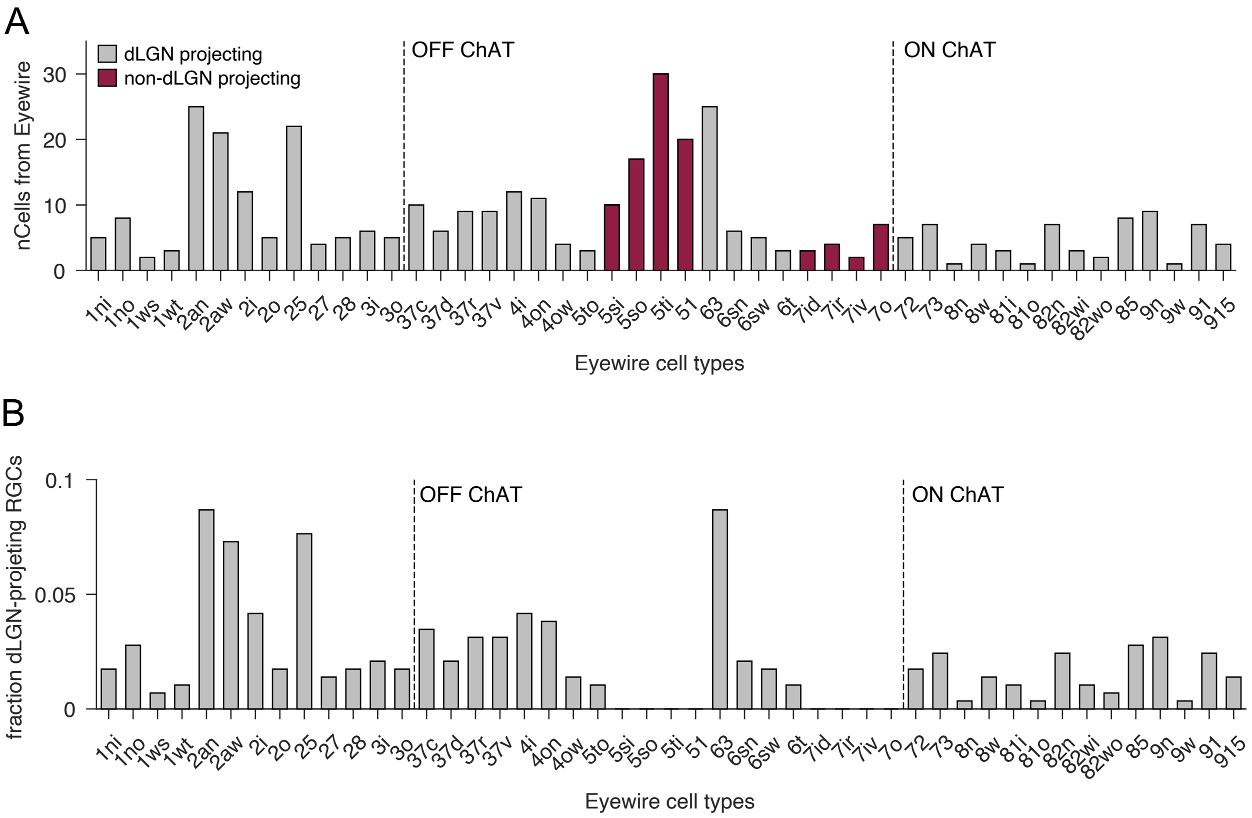


Supplementary Figure 4: (A) Distribution of numbers of cells of each type classified in the Eyewire dataset (Bae et al. 2018). Cell types highlighted in red are RGCs that do not project to dLGN (Kerschensteiner 2022). (B) Fraction of all dLGN-projecting RGCs by Eyewire cell type.
